# Gluebodies improve crystal reliability and diversity through transferable nanobody mutations that introduce constitutive close contacts

**DOI:** 10.1101/2022.07.26.501559

**Authors:** Mingda Ye, Mpho Makola, Joseph A. Newman, Michael Fairhead, Elizabeth Maclean, Nathan D. Wright, Lizbé Koekemoer, Andrew Thompson, Gustavo A. Bezerra, Gangshun Yi, Huanyu Li, Victor L. Rangel, Dimitrios Mamalis, Hazel Aitkenhead, Benjamin G. Davis, Robert J.C. Gilbert, Katharina Duerr, Opher Gileadi, Frank von Delft

**Affiliations:** Centre for Medicines Discovery, Nuffield Department of Medicine, University of Oxford, Oxford, UK; Division of Structural Biology, Wellcome Centre for Human Genetics, University of Oxford, Roosevelt Drive, Oxford OX3 7BN, UK; Present address: The Walter and Eliza Hall Institute of Medical Research, 1G, Royal Parade, Parkville, Victoria, 3052, Australia; Present address: Bicycle Therapeutics Plc, Cambridge, UK; Laboratory of Protein Crystallography, School of Pharmaceutical Sciences of Ribeirão Preto, University of São Paulo, Ribeirão Preto, São Paulo, Brazil; Present address: Evotec Ltd, Oxford, UK; Department of Chemistry, University of Oxford, Oxford, UK; The Rosalind Franklin Institute, Oxfordshire, Oxford, UK; Department of Pharmacology, University of Oxford, Oxford, UK; Calleva Research Centre for Evolution and Human Sciences, Magdalen College, University of Oxford, Oxford OX1 4AU, UK; OMass Therapeutics, Ltd., Oxford, UK; SGC Karolinska Center for Molecular Medicine, Karolinska University Hospital, 171 76 Stockholm, Sweden; Diamond Light Source, Harwell Science and Innovation Campus, Didcot, UK; Research Complex at Harwell, Harwell Science and Innovation Campus, Didcot, UK; Department of Biochemistry, University of Johannesburg, Auckland Park, 2006, South Africa

## Abstract

The design of proteins that may assemble in a manner that is transferable and modular remains an enduring challenge. In particular, obtaining well-diffracting protein crystals suitable for characterizing ligands or drug candidates and understanding different protein conformations remains a bottleneck for structural studies. Using nanobodies as crystallization chaperones is one strategy to address the problem, but its reliability is uncharacterized and, in this study, we observed it to have a limited success rate. Here we show that by exploring and testing the nanobody-nanobody interfaces predominant in >200 combinations of surface mutations in multiple iterations we can engineer robust crystallization behaviour into the nanobody scaffold. Strikingly, this survey yielded multiple polymorphs, all mediated by the same interface. The resulting ‘Gluebodies’ (Gbs) provide far superior resolution and reliability of diffraction and can be routinely generated for chaperone experiments. We furthermore show that Gbs cannot rescue intrinsically non-crystallizing proteins, but instead are a powerful approach to improve the packing and resolution limit of poorly diffracting crystals. The discovery of an engineered, preferred nanobody interface that arises under kinetic control - trapped here by irreversible crystallization - embodies a protein assembly strategy that could prove even more broadly useful for modular assembly trapped by other irreversible methods.

## Introduction

Protein X-ray crystallography has long been a routinely used method to elucidate the near-atomic structure of proteins of interest. However, enabling the protein to consistently crystallize can prove highly troublesome, with no substantial methodological breakthroughs for many decades. The mechanistic specifics remain poorly understood, so that there is no systematic route to coax a protein to pack in an ordered lattice ^1–5^. Nevertheless, numerous techniques have emerged throughout the past decade to aid crystallization and they generally follow one of four general strategies.

The first is to vary the protein environment by exploring crystallization solution, temperature or physical format ^6, 7^. This approach has been thoroughly commercialized for over two decades, with a large repertoire of crystallization screens purchasable from vendors, and many laboratories equipped with various systems of automation ^8^.

The second strategy is to modify the protein itself to favour crystallization, either by introducing major changes to the protein through varying the expression construct ^9^, or by subtle changes on the protein surface, such as by surface entropy reduction ^10^, chemical modifications ^11^ or the ‘crystal epitope’ approach ^12, 13^. Using protein orthologues that crystallize more easily is also an alternative ^14^.

The third approach is to introduce a natural partner that forms a complex with the target of interest. The natural binding partner, including ligands and proteins, could potentially stabilize the target or introduce additional surface for forming crystal contacts ^15–18^.

The fourth approach is to use a protein ‘chaperone’ engineered to provide the target with additional or alternative potential crystal contacts. Reported chaperones include fusion tags such as lysozyme ^19^ or BRIL ^20^; and protein binders such as Fabs ^21^, scFvs ^22^, nanobodies/sybodies ^23, 24^ or DARPins ^25^. The required use of fusion tags can be challenging since their addition disrupts the native structure, decreases protein yields or otherwise impedes biochemical behaviour.^9^ As a result, this strategy often requires extensive testing of a combinatorial matrix of constructs to identify the insertion point that optimizes protein packing and thus improves diffraction^.26^ Consequently, binder-assisted protein crystallization has tended to be used as a last resort for the most challenging targets, such as membrane proteins ^27^, because these tend to be long-term projects where the necessarily extended timescales required for obtaining binders are not seen as rate-limiting. Nevertheless, thanks to the improvement and diversification of binder selection methods, the generation of binders to assist structural studies is now increasingly attempted in many labs.^24, 28, 29^

Beyond these, more speculative suggestions to aid crystallization have also been reported, but have either not been well-developed (e.g. embedding the protein of interest into a highly porous crystal lattice ^30–32^), or have not yet, despite their ingenuity, achieved widespread usage and thus validation (e.g. imprinted polymers ^33^, microgravity ^34^, racemic protein crystallography ^35^).

Therefore, whilst none of these methods routinely allow enhancement of crystal reliability and diversity yet, binder-assisted crystallization perhaps holds the greatest promise to address several key crystallization challenges beyond simply obtaining a first crystal structure ^27^. Firstly, such binders mask out surface heterogeneity and reduce the entropy for forming stable crystal lattices ^36^. Secondly, they can lock the target protein into conformations that cannot be otherwise isolated ^37, 38^. Thirdly, they can help find considerably more robust crystallization systems that are required for structure-based lead discovery, including crystal-based fragment screening, since the introduction of new or additional crystal-crystal contacts mediated by the chaperone can systematically increase polymorphism ^39, 40^. Finally, by engineering binder scaffolds, symmetry can be introduced to the binder:target complex to further facilitate crystallization processes ^41, 42^.

Significant challenges prevent current realization of this promise. Although benefits have been suggested in human Fabs (e.g. by shortening the FG loop by two residues)^43^ or diabody scaffolds (e.g. at the V_H_-V_H_ interface)^44^ these remain bespoke and non-modular. Moreover, binders such as Fabs and diabodies are relatively difficult to express and purify, rendering the engineering process arduous.

Here, we demonstrate a workflow in the readily-manipulated nanobody scaffold where, firstly, a wholly crystallization-ineffective nanobody can be made effective by a limited number of mutations on the nanobody scaffold and, secondly, enhanced with mutational screening of more than 200 constructs designed through data mining. This identifies four key mutations on the nanobody scaffold far from the CDR surfaces that lead to extensive crystal polymorphism and robust diffraction (Figure 1). Finally, we describe the ability of these engineered nanobodies to help enhance crystallizability of proteins reluctant to crystallize. This engineerable process, driven by an interface trapped under kinetic (here crystallization) control, should prove transferable not only to other targets but indeed to other widely-used chaperone scaffolds and even other kinetically-trapped processes.

**Figure 1.**
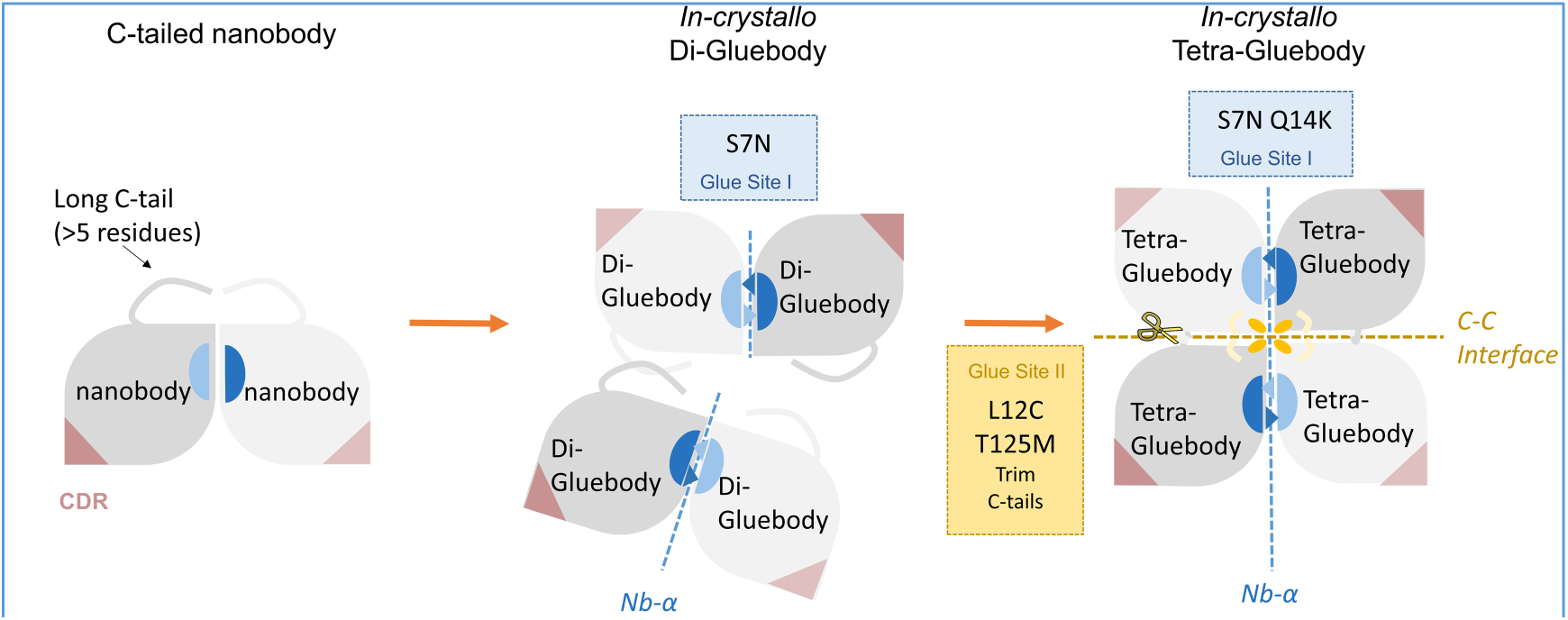
The Evolution of Nanobody scaffold to Gluebodies. Progressive sets of mutations that engineer the Gluebody interface lead to diverse polymorphs with improved crystallizability and diffraction quality. The putative crystallization patterns from the three scaffolds are presented from left to right with the scaffold type at the top of the sketches. The leaf shapes colored in grey and light grey represent nanobody molecules. CDRs are indicated in light brown. The Nb-α contacts are shown in blue and light blue patches. The long or short overhangs on the leaf shapes represent the C-terminal tails of the nanobody molecules. The orange ovals, blue triangles and pale hooks represent the key mutations on the nanobody scaffold. (C12, N7 & K14 and M125, respectively) The dashed lines represent nanobody-nanobody interfaces occurring in the putative crystallization patterns.

## Results

### Naïve nanobodies are unsatisfactory crystallization chaperones

We selected 17 nanobody-target pairs across 5 different target proteins (Extended Data Table 1 and Extended Data Figure 1). Among the targets, solved crystal structures (of apo RECQL5, RECQL5 bound with ADP/Mg ^45^ and apo WRN ^46^) were included as benchmark comparators. We performed standard crystallization trials of the 10 protein complexes (7 failed during purification) against two commercially-available coarse screens (192 conditions, 2 temperatures) for each protein complex. This yielded crystals for only one protein complex (MAGEB1), which diffracted well and could be readily solved (PDB ID 6R7T). This initial screen suggested that the naïve effectiveness of appropriately binding nanobodies as crystallization chaperones without engineering is less than 10%, demonstrating the need for dramatic improvement of the approach.

### A single nanobody surface mutation by the Crystal Epitope approach yields a first effective chaperone for RECQL5

The strategy of Crystal Epitope ^13, 47^ entails identifying short sequence motifs (3-6 residues) that frequently appear in crystal contacts in the Protein Data Bank (PDB). We have previously successfully used this strategy to obtain crystals of several different target proteins (e.g., BRD1A: 5AMF, GALT: 6GQD, Glycogenin-2 catalytic domain: 4UEG), and similarly we applied it to the non-CDRs of a specific nanobody targeting RECQL5.

74 crystal epitope mutations (variant class G0 and variant G1-001) were designed through 3 iterations on the scaffold (non-CDR, non-variable region) of a RECQL5 nanobody (Extended Data Figure 3, 3A and 3B, Extended Data Table 2). Purified nanobodies all co-migrated with RECQL5 on size exclusion chromatography, indicating adequate affinity of the nanobodies. The purified protein complexes were put into crystallization trials using the Hampton Index coarse screen, and 5 of the 74 yielded crystals (Extended Data Figure 3C, 3D and 3E). These variants shared one common mutation, D69Y, the single mutation of variant G1-001 (Extended Data Figure 2 and Extended Data Table 4) and crystals of the G1-001 complex diffracted to the highest resolution (2.76 Å). Consistent with this initial design, in the mutant structure, Y69 on the nanobody forms a hydrogen bond with D400 of RECQL5, which most likely stabilizes the crystal formation (Figure 6A). The striking effect that this emergent single mutation has on its crystallizability gave first proof-of-principle that the nanobody scaffold could be engineered for improved crystallization – a general screening approach can be applied to nanobodies targeting different proteins, thereby bootstrapping the chances to obtain a first crystal.

### A single nanobody-nanobody interface predominates in PDB structures

This demonstrated utility of non-CDR surface mutations increased crystallization propensity while preserving nanobody affinity for its target protein. Nonetheless, the effective mutations were not mediating nanobody-nanobody crystal contacts likely to build a modular lattice, and thus were only useful for their specific target RECQL5. To optimize a nanobody interface independent of target proteins, we next applied a general strategy applicable to all nanobody-protein complexes, by engineering only nanobody-nanobody interactions (Figure 2).

**Figure 2.**
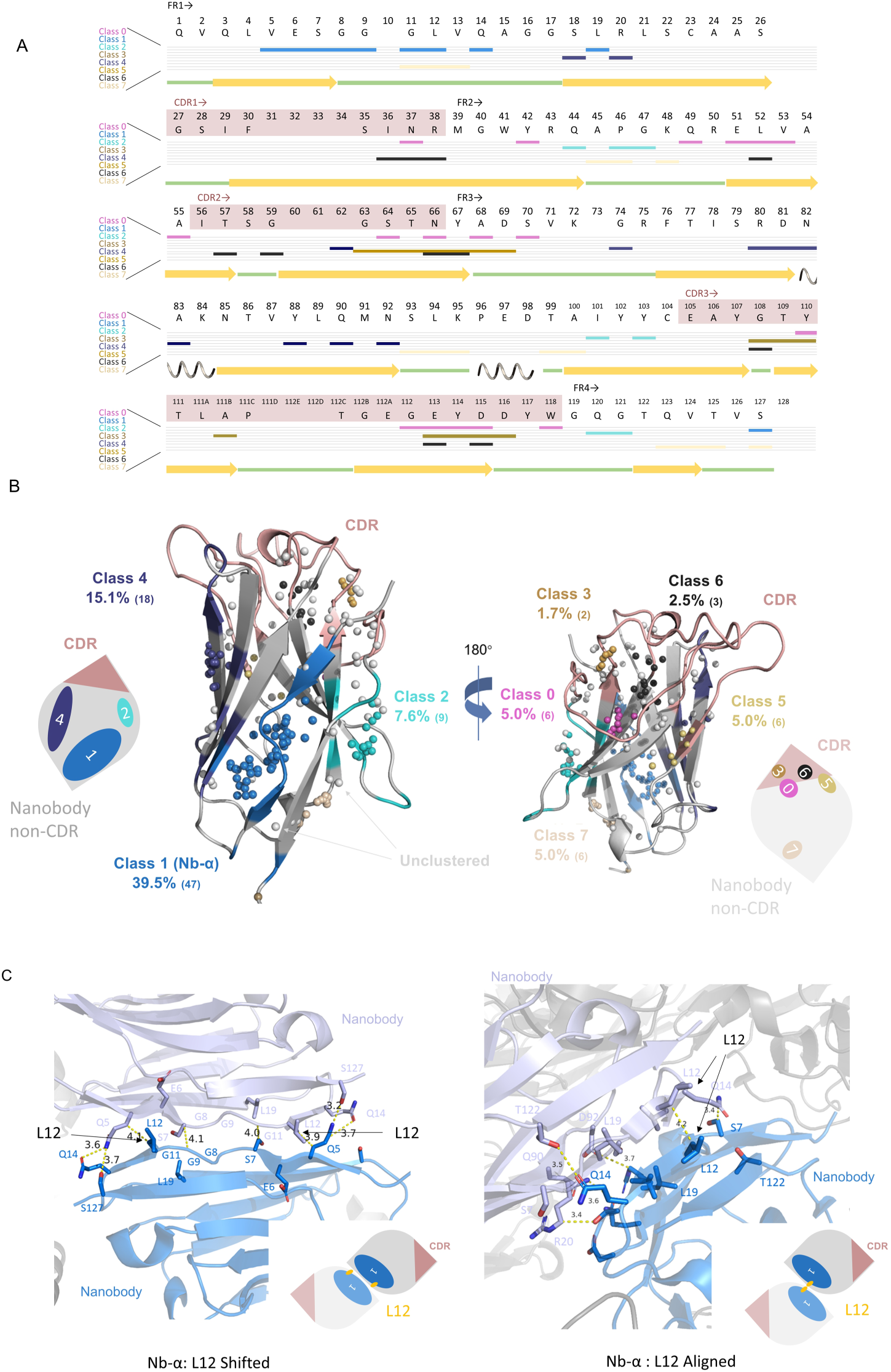
Nanobody-nanobody crystal contacts fall into limited classes. (A) Standard nanobody residue numbering of C-terminally tagged wild type RECQL5 nanobody (C-tag:WT) using an IMGT scheme provided by ANARCI ^49^. CDRs are in light brown background and non-CDRs are in white background. Participating residues of each category are indicated by a bar below the one-letter code. (B) DBSCAN clustering result: global view of major nanobody-nanobody crystal interfaces on a model drawn from chain E of 5O0W in PDB.

To understand how nanobodies commonly pack with each other in crystals, we examined the 335 nanobody-containing crystal structures in the PDB. We defined two molecules as forming crystal contacts if their nearest atoms are below 4 Å apart ^48^, and thus found 273 structures with nanobody-nanobody contacts. Contacts with more than 5 interacting residues were deemed suitable for engineering. In this way, interactions were refined to 356 large contacts across 119 structures for further analyses.

Through use of a bespoke Python script (using the Python interface in Pymol 2.7), we were able to extract all interface information in the 335 crystal structures and focused on the interfaces that only involved nanobodies. The nanobody:nanobody interfaces were represented by index arrays of the residues involved. The index array was then renumbered (using an IMGT scheme on the online server ANARCI ^49^, Figure 2A), enabling comparisons across the nanobodies appearing in different deposited structures. The renumbered arrays were further converted into coordinate matrices, which then immediately shrank into a 4-dimensional array (n*, μ_x_, μ_y_, μ_z_) (Extended Data Figure 4). The four items in the array represent 1/3 of the number of residues in the interface, and the mean values of the X, Y, Z coordinates of the residues in the interface, respectively. Selected nanobody interfaces were then mapped onto a single nanobody structure (PDB id 5O0W) using their coordinate information (Extended Data Figure 4). As numeric representations of the interfaces, 4-dimensional vectors were used and we further performed density-based spatial clustering (DBSCAN) on the 356 vectors (Extended Data Figure 4).

Clustering revealed eight classes of nanobody-nanobody crystal contacts. Four of them (Class 1, 2, 4, 7) away from the CDRs, account for 39.5%, 7.6%, 15.1% and 5.0% of the total (Figure 2B). The most prevalent, termed ‘Nb-α’ (39.5%), is an edge-to-edge Class 1 interaction of the two beta sheets at the N terminus (Figure 2C). The Nb-α contact typically has 10 residues interacting in this type of contact, creating a plethora of options for engineering an interface that might be more likely to crystallize.

The other contact classes 0, 3, 5 and 6 are located in or very close to CDR regions, and therefore are not likely to mediate crystal packing and target binding at the same time. Contact classes 2 and 7 have relatively rare appearances in PDB. Finally, class 4 contact also presented a potentially good surface for engineering; however, it appeared with half the frequency and is closer to CDRs. We therefore chose to focus on surface engineering within the Nb-α contact family.

### Iterations of nanobody scaffold mutagenesis yield consistently diffracting crystals

Based on these mining results and within our RECQL5:nanobody system, we designed five generations of nanobody scaffolds (G1-G5) of more than 200 constructs with different combinations of Nb-α mutations and crystal epitope mutations (Extended Data Figure 2 and Figure 3A). This used a rationale for mutational design around Nb-α of interactions that might enhance interface affinity (e.g. salt bridges, hydrogen bonds and potential disulfide bonds).

**Figure 3.**
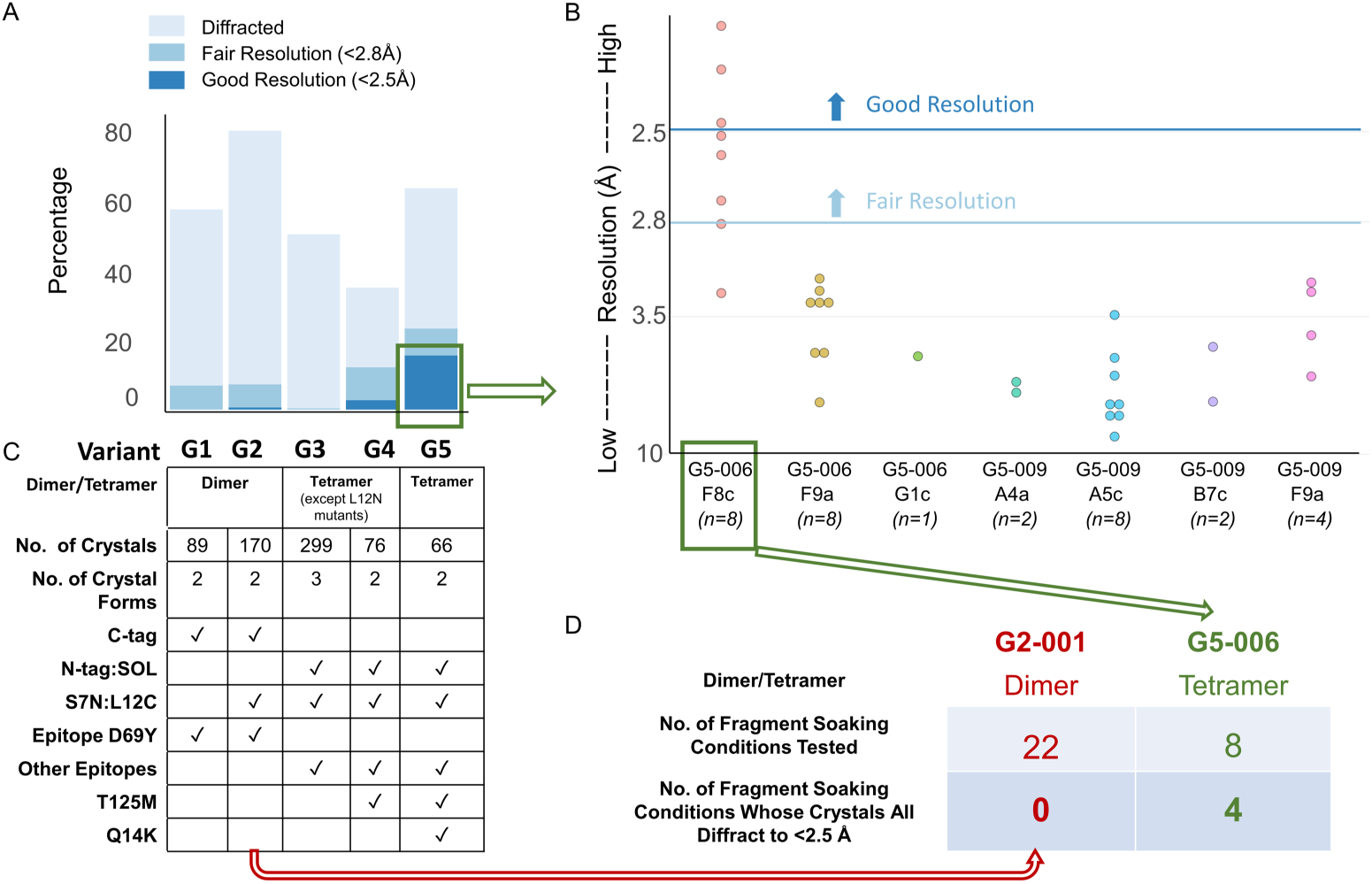
Consistently well-diffracting crystal emerge only after several iterations of mutagenesis around Class 1 contact Nb-α. (A) Diffraction quality of different groups of variants measured by the percentage of crystals which diffract to high resolution. The numbers of crystals and mutations present in each group of variants are indicated in the table below each bar. (B) Diffraction quality of different nanobody variant and condition pairs of RECQL5:nanobody variant complexes presented as a dotplot. (C) Summary of characters of G1-G5 variants, number of crystals tested and number of crystal forms observed. (D) A table summary of diffraction after testing cryoprotectants against crystals of G2-001 and G5-006 (details in Extended Data Figure 8).

This yielded highly effective reagents; retrospective analysis allows extraction of the mutational origins underpinning these successful iterations. In brief, G1 and G2 mutants, derived from the original RECQL5 nanobody scaffold, bore a C-terminal 6xHis tag that is cleaved during purification (resulting in 6-residue tail ENLYFQ after TEV protease-mediated cleavage). G2 crystals with the S7N:L12C mutations showed higher diffraction rate, however the high-resolution fraction did not significantly increase. Next, in G3, solubility mutations (G40T:Q49E:L52W:I101V, abbreviated SOL, that mimic the well-expressing GFP-enhancer nanobody from PDB:3K1K) were combined with both movement of the C-terminal 6xHis tag to an N-terminal MBP-6xHis tag (which does not yield an ENLYFQ) as well as epitope mutations beyond D69Y. Although G3 crystals showed initially worsened diffraction quality, two mutations (T125M and Q14K) based on G3 scaffolds, drove significantly improved diffraction quality in G5.

Each colored sphere on the structure represents the average location of participating residues of an interface analyzed. Grey spheres represent ungrouped nanobody interfaces in the analysis. The three largest interface patterns are colored on the cartoon of the nanobody structure with respective class colors. Simplified nanobody sketches are shown at the corners with CDRs (light brown) and numbered contact classes, indicating their relative locations on a nanobody. The areas of contact color patches are proportional to their respective percentages. Colours are consistent in A and B. (C) Typical close views of Class 1 (Nb-α) Nanobody-nanobody crystal contacts. L12 Shifted type is shown in the left panel, and L12 Aligned type is shown in the right panel. Structure cartoons in marine and light blue represent two nanobody molecules participating in the interface. Residues represented as sticks are interacting residues. The simplified nanobody sketches are shown at the bottom corners (See Extended Data Figure 4 and Extended Data Table 2 for detailed information of each class of nanobody-nanobody crystal contacts)

We explored in further detail the crystals derived from G5. There were several crystallization conditions yielding crystals within just 2 days; these were selected. 10-12 crystals from each variant-condition pair were harvested for diffraction analysis. The pair G5-006-F8c had a significantly larger fraction of crystals diffracting better than 2.5 Å (Figure 3B). Improvements were also importantly obtained in processes employing optimization of cryo-protectant and DMSO concentrations (i.e. under potential fragment-soaking conditions). Out of the 8 conditions we tested, 100% crystals from 4 conditions diffracted better than 2.5 Å (Figure 3C and Extended Data Figure 8); this is a critically desirable resolution threshold to reliably determine the ligand-binding conformations^50^. Notably, when comparing G5-006 with G2-001, the diffraction quality of G5-006 generation-5 crystals was significantly improved over generation-2, consistent with successful iterative engineering.

### Constitutive nanobody interfaces were present in 6 different crystal forms

In the five generations of nanobody scaffolds, we looked for constitutive nanobody-nanobody interfaces that might be independent of target protein, consistent with our design hypothesis.

Starting with Nb-α in G2, upon combining S7N:L12C with prior G1, the crystallization propensity of RECQL5:nanobody complex increased dramatically for both N-terminally (N-tag) and C-terminally (C-tag) tagged variants (Figure 4A and 4B). Comparison of the structures of G1-001 with G2-001 reveals an additional hydrogen bond in Nb-α introduced by S7N between the side chain N7 and the carbonyl oxygen of A15 (Figure 4C). These S7N:L12C mutations were retained in the following iterations of mutagenesis.

**Figure 4.**
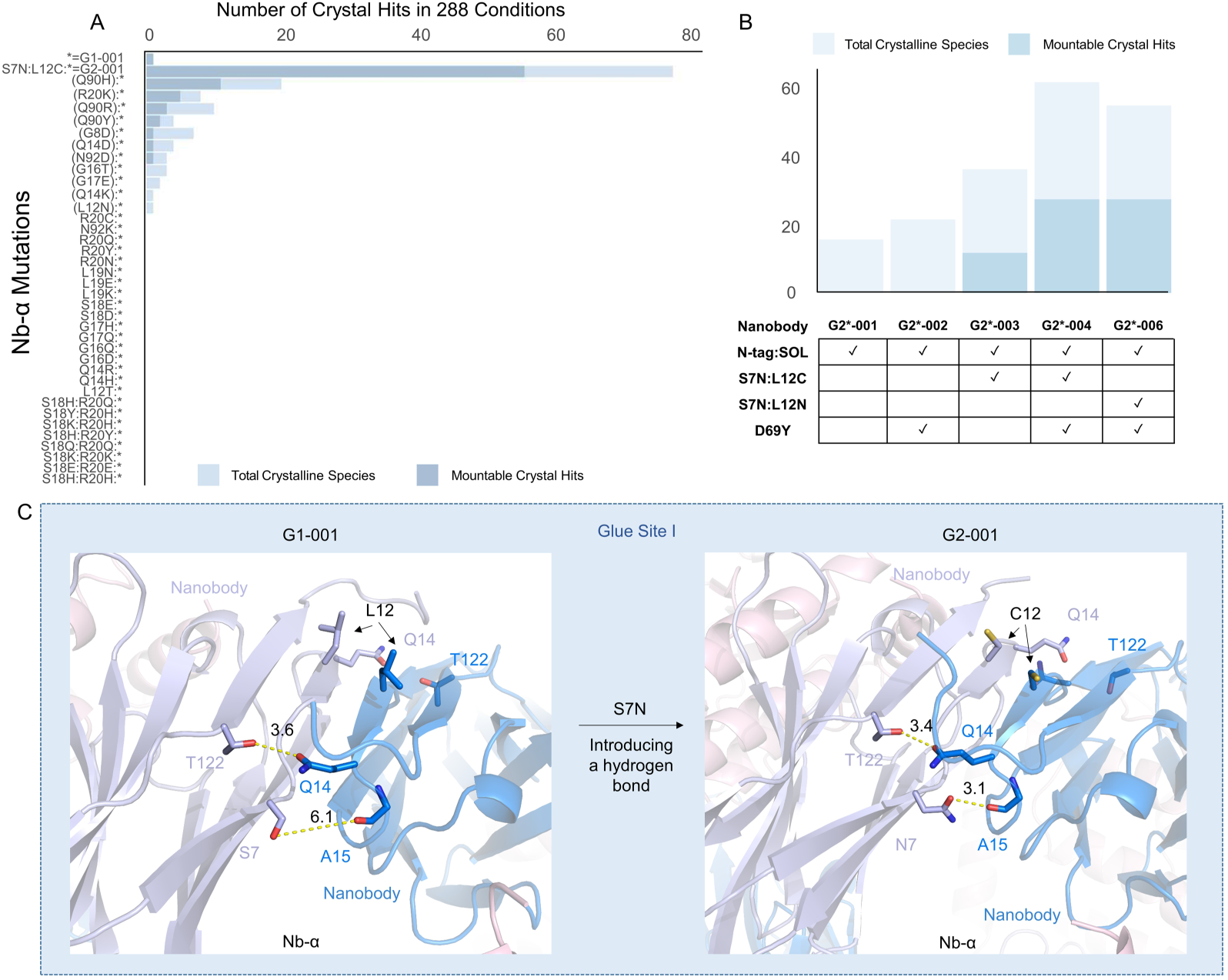
Mutation S7N enhances Nb-α by introducing an additional pair of hydrogen bonds. (A) Hampton Index 3 (HIN3) triplicate drop coarse crystallization screen experiments show that S7N:L12C stands out among Nb-α mutations in significantly increasing the crystallization propensity of the RECQL5:nanobody complex. The asterisk represents the variant complex G1-001. (B) HIN3 triplicate drop coarse crystallization screen experiments show that S7N:L12C significantly increases the crystallization propensity of the N-tag:SOL variant of RECQL5:nanobody complex with or without D69Y as the crystal epitope mutation. (C) Close views of the Glue Site I before and after the mutation. S7N (stick representation) introduces an additional hydrogen bond with the carbonyl oxygen of A15 (stick representation) on the opposite nanobody molecule. Oxygen atoms are colored red and Nitrogen atoms are colored blue. The distances between the atoms connected by dash yellow lines are shown in Å.

Different combinations of crystal epitope mutations and Nb-α mutations were further explored in G3. We replaced D69Y with 9 other crystal epitope mutations in G3 and changed the terminal tag position (see above). Crystallization patterns changed dramatically with most G3 crystals requiring high salt conditions whilst most G2 crystals required PEG conditions (Figure 5A). Notably, Cys12 appeared important in the crystallization of G3 – introduction of the C12N mutation decreased the average number of observed hits significantly (Figure 5A). Moreover, mutation of Cys12 to other residues (including via chemical mutation to dehydroalanine Dha^51^, Extended Data Figure 7) resulted in even ‘good crystallizers’ immediately losing crystallizing ability (Figure 5B).

**Figure 5.**
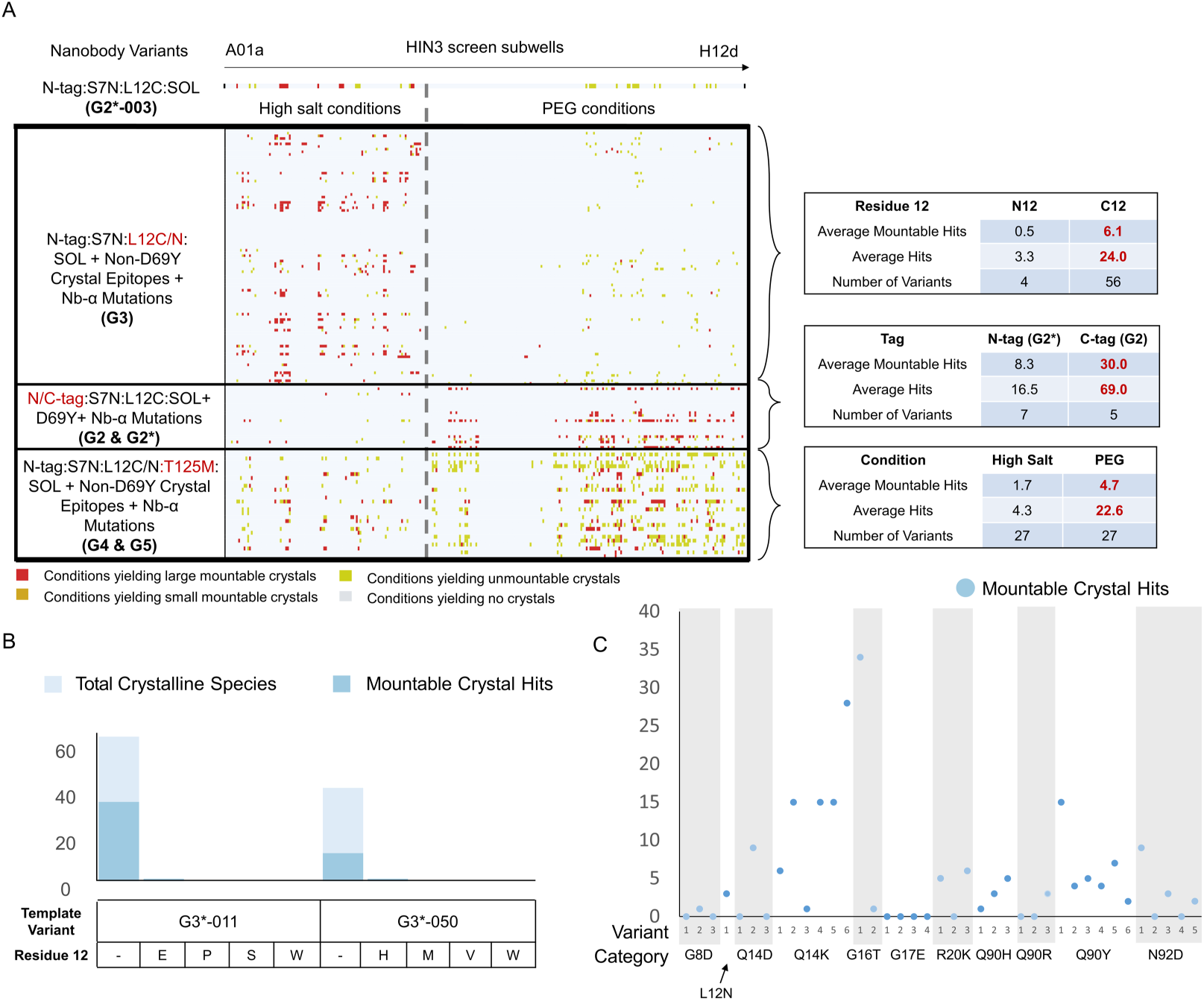
G2, G3 and G4&5 have different crystallization patterns in the Hampton Index coarse screen. (A) Hampton Index 3 coarse screen triplicate crystal trials of RECQL5:nanobody complex with various mutations on the nanobody scaffold. Each row represents a nanobody construct, and each column in the chart represents a crystallization condition in the Hampton Index 3 coarse screen. The conditions are linearized sequentially from A01a on the left to H12d on the right. To the left of the grey dashed line are high salt conditions and to the right are PEG conditions. The color of each cell in the row represents whether there are crystals growing in that condition and the quality of the crystals. Some statistics of each group of nanobody variants are generalized on the right of the main plot. See Extended Data Table 5 for detailed information of what each row and column represents. (B) The RECQL5 nanobody loses crystallization propensity when C12 is mutated to a selection of other residues. (C) HIN3 triplicate drop coarse crystallization screen experiments show Q14K led to higher mountable crystal hits among a set of Nb-α mutations in G4&G5. The graph shows mountable crystal counts against each variant in respective Nb-α mutation categories.

Since Cys12 was demonstrated to be important in improving crystallization, the area around this striking focal point lynchpin residue was futher explored by mutagenesis of neighbouring residues. In particular, when T125M was introduced in G4 and G5, the crystallization pattern again changed, expanding hits under both high salt and PEG conditions (Figure 5A). Furthermore, with the Nb-α mutation Q14K in G5, the crystallization propensity was further improved (Figure 5C) with even noticeable growth of large crystals. It should be noted that ambiguity in the electron densities captured from the K14 side chains means that the mechanistic origin of this improvement remain unknown, further highlighting the value in discovery of our combined, iterative computation-plus-empirical approach over one based on structural analyses alone.

Interestingly, the significant crystallization pattern change observed on moving from G2 to G3 could also be explained. Consistent with a stepped evolutionary approach to these hierarchical changes, an entirely different nanobody arrangement emerges *in crystallo*. Instead of a dimer nanobody core (Figure 6A left panel) mediated by Nb-α in G1 and G2, we instead observed a tetrameric nanobody core (Figure 6A right panel) in G3. In this tetrameric nanobody arrangement, the Cys12 residues are in the core of the interface in contact with each other. We named this emergent interface, which perpendicular to Nb-α, the ‘C-C interface’. The emergence of the novel C-C interface at G3 can be further understood. Due to the geometric arrangement of the four nanobody molecules, the tetrameric core in G3 would not form with the 6-residue C-terminal tail ENLYFQ still present; this tail can be observed, for example, in the G2-001 structure ‘hanging over’ the critical Cys12 residues (Figure 6A left panel). The critical role of Cys12 can also be understood in forming the tetrameric nanobody arrangement. When mutated to Asn12, the C-C interface dissociates and only the dimeric nanobody arrangement is observed (Figure 6B), resulting in a much lower resolution (Extended Data Table 4).

**Figure 6.**
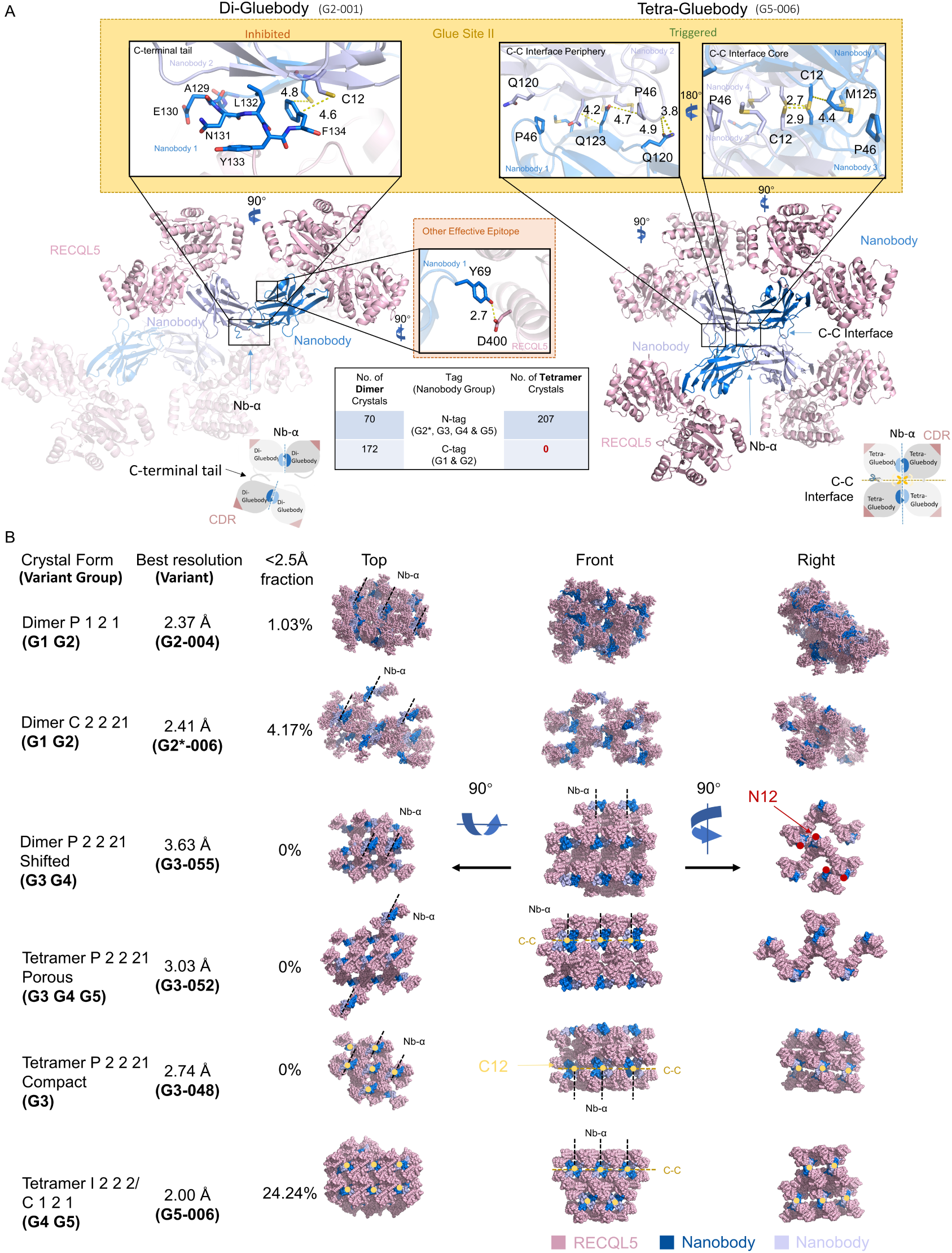
Two constitutive nanobody interfaces are found in 6 different crystal forms in Gluebodies. (A) *In-crystallo* dimeric (Di-Gluebody) and *in-crystallo* tetrameric (Tetra-Gluebody) crystallization pattern of RECQL5:nanobody with close views of Glue Site II and the crystal epitope site of Y69. The left panel shows RECQL5:nanobody complex crystallizes in a dimeric pattern when mutation D69Y and the C-terminal tail exist on the nanobody scaffold. The cartoon structure with high transparency is a symmetry mate of the dimer. The schematic of the dimer pattern is shown on the bottom left. The right panel shows the tetrameric crystallization. The M125 has a longer side chain than the T125 and occupies the cavity around the cysteine core within the C-C interface. The schematic of the tetramer pattern is shown on the bottom right. Oxygen atoms are colored red and Nitrogen atoms are colored blue. The distances between the atoms connected by dash yellow lines are shown in Å. (B) Six different crystal forms of the RECQL5:nanobody complex with respective mutations on the nanobody scaffold are presented in three perspectives (top, front and right views). RECQL5 molecules are in light-pink, and nanobody molecules are in marine and light-blue. C12 residues are presented as yellow dots on the structures and N12 residues are presented as red dots. Nb-α and C-C interfaces are indicated by dashed lines in the lattices. Resolution indicated here uses the criteria CC ½ > 0.3 from the ISPYB auto-processing pipeline without any further data truncation.

From G1-G3 we already observed five different crystal forms (Figure 6B). Dimer P 1 2 1 and Dimer C 2 2 21 existed mostly in PEG conditions. Tetramer P 2 2 21 Porous/Compact and Dimer P 2 2 21 shifted mostly existed in high salt condition. In G4 and G5 we observed crystals in both high salt and PEG conditions. The high salt crystals of G4&5 were Tetramer P 2 2 21 Porous form, which we already observed in G3. The reason why G4 and G5 crystallized under drastically different conditions might be that it unlocked an additional crystal form Tetramer I 2 2 2/C1 2 1. Notably, this crystal form led to our best-diffracting data (Extended Data Figure 5), indicating that crystal packing was also improved by the mutation T125M.

Among all six crystal forms, the Nb-α interface was present, and among all tetramer crystal forms, the C-C interface was present (Figure 6B). This indicated that these two interfaces are key, mediating crystal packing under certain conditions like the glue-ing of surfaces. We therefore termed the nanobody scaffolds with our modifications that enhance these two interfaces as ‘Gluebodies’ (Gbs). We also termed interacting residues in Nb-α as ‘Glue site I’ (Figure 4C and 6A) and interacting residues in the C-C interface as ‘Glue site II’ (Figure 6A).

### Gluebodies can improve crystallizing proteins but do not rescue non-crystallizing proteins

The sets of mutations around the Nb-α interface on the Gluebody are demonstrated to improve the crystallization propensity of the nanobody for the model protein RECQL5, by enabling various crystal forms via dimeric or tetrameric Gluebody arrangements. However, the improvement of the nanobody alone was not sufficient, in our hands, to yield crystals for non-crystallizing proteins. This was attempted on five other proteins (Table 1 and Extended Data Figure 6), but yielded only poor or problematic crystals. Only 6-8 Å diffraction was obtained from TUT4 and PRNP crystals, and for a Major Facilitator Superfamily (MFS) protein, the crystals of the Gluebody complex did not diffract as well as the original nanobody scaffold. Thus, the associating interfaces of Gluebodies provided initial hits for other targets reluctant to crystallize, but given the low-resolution diffraction we observed, more target construct screening or testing more crystallization conditions are probably necessary on a case-by-case basis.

**Table 1.** Experimental details of the crystallization trials of several targets in complex with respective Gluebodies. The WRN:Gluebody crystal diffracted to 2 Å, but the phasing failed, resulting in an unsolvable dataset.

| *Target* | *No. of Glubodies tested* | *No. of crystal plates with Gluebodies* | *Lipid Cubic Phase?* | *Crystals with Gluebody* | *Crystals with original nanobody* | *Comments* |
| --- | --- | --- | --- | --- | --- | --- |
| **TUT4** | 5 | 20 | No | 6-8 Å | None |  |
| **WRN** | 8 | 18 | No | 2 Å | None | Single crystal, unreproducible, phasing failed |
| **PRNP** | 1 | 2 | No | 6-8 Å | None |  |
| **MFS protein** | 2 | 12 | Yes | No diffraction | 7-10 Å |  |
| **MAGEC2** | 3 | 6 | No | None | None |  |

## Discussion

In this study, we identified multiple crystal polymorphs in a model (RECQL5) protein target:nanobody system through iterative engineering of the nanobody scaffold (Extended Data Figure 5). This Gluebody method provides a route to novel and better behaving crystal polymorphs. Strikingly, the polymorphs are excitingly diverse, including a novel interface-driven tetramer lattice, despite involving only a single target protein. This contrasts strongly with the polymorphic behaviours of naïve nanobody scaffolds, as inferred from the PDB, where, for instance, the tetrameric nanobody pattern is present only twice (PDB IDs 6OS1 and 6OS2), notably with fewer interacting residues than in our Gluebody.

This engineering workflow was importantly enabled by state-of-the-art technologies (e.g. high-throughput protein purification and crystal harvesting, X-ray diffraction in large synchrotrons) that vitally streamlined iterative analyses of the hundreds of mutant crystal growth and diffractions that were required. In this way, we show that it has now become viable to monitor ‘phenotypic’ changes of crystallization patterns and so utilize X-ray crystallography as a powerful (and here the sole) methodology for iterative protein engineering.

For such ‘trapping’, kinetics (here of crystallization) plays a pivotal role. Therefore, other biophysical methods (such as BLI, MST and ITC) that may measure binding affinities and so provide insight into thermodynamics are neither direct nor necessarily relevant outputs. Here this is vitally relevant when trapping interfaces for better crystal packing, involving transient and dynamic interactions within one type of protein molecule, and with only subtle changes in affinities when mutations are applied. In this way, X-ray crystallography has allowed unique discovery of a suggested kinetically trappable interface (see below). It also suggests that striving for affinity alone (i.e. thermodynamic measures) in chaperones is not necessarily the correct design approach.

Key mechanistic observations arise from the analysis of evolution of this engineering workflow that reveal how interfaces may be ‘glued’ through kinetic trapping (here provided by crystallization). First, by introducing S7N Glue Site I mutations (Figure 4C, 6A and 1) crystallization is driven by the *in* situ formation of Gb dimers. Second, our data reveal a key role for the Gb C-terminus as an emergent interface (Glue site II) that drives polymorphism. Analyses reveal that our discovery is consistent with missing interfaces in current structures; the PDB reveals a trend of C-terminal tags attached (63% out of 335 crystal structures in PDB have more than 5 residues at their C-terminal sequence). As we explicitly show here, such ‘overhang’ residues hinder both the formation of the Gb C-C interface and impede the tetrameric assembly form by occluding C12. The combination of free C-terminal tail, Q14K and Glue Site II mutations (L12C, T125M) (Figure 6A and 1) trigger tetrameric crystallization polymorphs. This arises through a combination of glue sites as trapping interfaces: dimerization of diGb gives tetraGb *in crystallo* (Figure 6B).

Notably, although tetramers are formed in this way, the revealed formation of Gb dimers (*in crystallo* ‘di-gluebodies’) via various modes suggests that other trapping modes of interfaces may be feasible. Foremost amongst these would be the translation of this *in crystallo* trapping to a mode of in solution trapping enabled perhaps by chemically-controlled covalent bond formation. Such chemically-linked dimeric Gluebody complexes would be a scaffold that might combine the advantages of Gbs but with added introduction of symmetry. More importantly, if sufficient rigidity were achieved, it would be a highly modular ‘plug-and-play’ tool without the need of protein fusions via molecular biology, and thus potentially provide a powerful tool to generate protein assemblies for small-protein cryo-EM structure determination that would provide advantages over existing and often complex monovalent tools (e.g. DARPin cages^52, 53^, the Megabodies^54^, Legobodies ^55^). Relevantly, in the tetrameric crystallization pattern we observed 4 Cys residues at the C-C interface core. Yet, despite clear electron density between the two sulfur atoms in each pair of Cys residues (Extended Data Figure 10), the distances between two sulfur atoms of around 2.8 Å preclude the existence of a disulfide bond (∼ 2.05 Å ^56^), at least *in crystallo*. Work towards this goal of in solution covalent kinetic trapping, uniquely derived from the work here of X-ray enabled interface engineering of Gluebodies that kinetically trap through crystallization is the subject of a following manuscript.

## Methods

### Crystal contact analysis of structures containing nanobodies in PDB

Crystal structures were fetched and analyzed using the open-source PYMOL package in Python ^57^. The workflow of analysis is described in results and shown in detail in Extended Data Figure 4. Finally, a few interfaces that fell into the noise category were manually inspected and added to respective categories resulting in the final classification.

### High-throughput cloning, expression and purification of nanobody variants

The nanobody constructs of G0-002 to G0-074 and G1-001 were generated by the site-directed mutagenesis method based on G0-001 using pNIC-CTH0 as the vector plasmid. The nanobody construct of G2-001 was generated by mutagenesis method based on G1-001 using pNIC-CTH0 as the vector plasmid. The nanobody constructs of G1-002 to G1-091 were generated by mutagenesis method based on G1-001 using pNIC-CTBH as the vector plasmid. The nanobody constructs of G2*-005 to G2*-016 were generated by mutagenesis method based on G2*-004 using pNIC-MBP2-LIC as the vector plasmid and G2-002 to G2-013 using pNIC-CTBH as the vector plasmid with the same method and template DNA. G3*-001 to G3*-019, G3*-020 to G3*-038, G3*-039 to G3*-057, G3*-058 to G3*-076 and G3*-077 to G3*-095 were generated using mutagenesis based on templates G3-048, G3-055, G3-052, G3*-011 and G3*-050 respectively. G3-001 to G3-096 and G2*-001 to G2*-004 were directly synthesized as DNA fragments and subcloned into pNIC-MBP2-LIC vectors. G4-001 to G4-087 and G5-001 to G5-009 were also based on the synthesized DNA fragments using a different set of primers in the PCR step. All cloning procedures followed ligation independent cloning described previously^9, 58^.

Constructs of nanobody variants were transformed into Single step (KRX) E.coli strain (Promega) at the end of molecular cloning. Single colony was inoculated into 1 ml of LB media (Merck) and incubated overnight. Starter culture of each nanobody was then inoculated into 100ml autoinduction media (Formedia) containing 0.1% Rhamnose (Sigma), 0.01% antifoam 204 (Sigma) and 0.05 mg/ml Kanamycin (Sigma), at 37°C for 5.5 hours followed by 40-44 hours at 18°C. The base buffer we used for purification contained 5% glycerol, 10 mM HEPES (pH 7.5), 500 mM NaCl, 0.5 mM TCEP. Bacteria for each nanobody were harvested at 4000 g and resuspended in 10ml base buffer + 30 mM imidazole, 1% Triton, 0.5 mg/ml lysozyme, 10 ug/ml homemade benzonase, followed by storage in -80°C freezer overnight for complete cell lysis. The purification started next day with thawing the frozen pallets in room temperature water bath, followed by centrifugation at 5000 g for 1 h to obtain clear supernatant. The supernatant for each nanobody was then applied to one 1 ml Ni-NTA pre-packed column (GE healthcare) pre-equilibrated with base buffer + 30 mM imidazole. After thoroughly washing the Ni-NTA columns with base buffer, nanobody was eluted using 2.5 ml of base buffer + 500 mM imidazole directly into base buffer equilibrated PD-10 columns (GE healthcare). 3.5 ml of base buffer was then applied to PD10 columns to elute the nanobody protein base buffer without imidazole. Then TEV protease was added to protein solution with a 1:10 mass ratio for overnight incubation, and 10 mM imidazole was also added in the solution. In the next day, two Ni-NTA columns were pre-equilibrated for each nanobody with base buffer + 10 mM imidazole and the nanobody solution with TEV was applied to the columns to get rid of nanobody with uncleaved tags, TEV protease and contaminants. Flow-through fractions were collected and 2 ml of base buffer was applied to the Ni-NTA columns to flush all nanobody through. The collected fractions were ready for complex preparation.

### Expression and Purification of RECQL5

The truncated RECQL5 protein (11-453) was subcloned into pNIC-Bsa4 vector with TEV cleavable 6xHIS tag at the N terminus. The protein was expressed using BL21-DE3-pRARE strain in auto-induction TB media (Formedia) containing Kanamycin and 0.01% antifoam 204 at 37°C for 5.5 hours followed by 40-44 hours at 18°C. The base buffer we used for purification contained 5% glycerol, 10 mM HEPES (pH 7.5), 500 mM NaCl, 0.5 mM TCEP. Bacteria were harvested by centrifugation at 4000 g and re-suspended in 3 times volume of base buffer + 30 mM Imidazole, 1% Triton, 0.5 mg/ml lysozyme, 10 ug/ml homemade benzonase, followed by storage in -80°C freezer overnight for complete cell lysis. The purification started next day with thawing the frozen pallets in room temperature water bath, followed by centrifugation at 5000 g for 1h to obtain clear supernatant. The supernatant was then applied to Ni-NTA pre-packed columns (GE healthcare) pre-equilibrated with base buffer + 30 mM imidazole. After thoroughly washing the Ni-NTA columns with base buffer, protein was eluted using 2.5 ml of base buffer + 500 mM imidazole directly into base buffer equilibrated PD-10 columns (GE healthcare). 3.5ml of base buffer was then applied to PD10 columns to elute the RECQL5 protein in base buffer. Then TEV protease was added to protein solution with a 1:10 mass ratio for overnight incubation, 20 mM imidazole was also added in the solution. In the next day, twice quantity of Ni-NTA columns were pre-equilibrated with base buffer + 20 mM imidazole and the RECQL5 solution with TEV was applied to the columns to get rid of RECQL5 with uncleaved 6xHIS tag, TEV protease and contaminants. Flow-through fractions were collected, and flash frozen for making nanobody complexes.

### Dehydroalanine formation

To a solution of G5-007 nanobody (1 mL, 1 mg/mL, 72 μM) in Na_2_HPO_4_ buffer (50 mM, pH 8.0), 0.28 mg (25 eq.) of DTT were added. The solution was shaken at room temperature for 20 min and then the protein was treated with 22 μL of 0.5M DBHDA (2,5-dibromohexanediamide) in DMSO (500 eq.) and heated to 37 °C for 4 h, at which point analysis by mass spectrometry showed reaction completion. The protein was purified using a PD Miditrap™ G-25 column (Cytiva #28918008) pre-equilibrated with Na_2_HPO_4_ buffer (50 mM, pH 8.0).

### Crystallization of RECQL5:nanobody complex variants and X-ray diffraction

Purified RECQL5 and nanobody were mixed at a ratio of 1:1.5 and concentrated using 10 kDa concentrator (Amicon and Corning), followed by Sepax SRT SEC-300 HPLC or Superdex-200 SEC (Cytiva). The peak fractions containing both RECQL5 and nanobody were pooled and concentrated to certain extent (Extended Data table 3) for crystallization trials. Crystal plates were set up by Mosquito (Model No. TC1100-1100, TTP Labtech) using Swiss-CI 3 drop plates with precipitants from Hampton Index screen (HIN3 HT-96, Molecular dimensions). Plates were sealed and sent into Formulatrix imager (Model No.R1-1000) and imaged at 12 hours, 1 day, 4 days, 7 days, 14 days, 28 days and 56 days. Crystals were harvested using Shifter^59^ with 17% glycerol for high salt conditions or ethylene glycol for PEG conditions added as cryo-protectant, then snap frozen and shipped to multiple beamlines at Diamond Light Source for X-ray diffraction screening and data collection.

### Structure determination of RECQL5:nanobody complex variants

Diffraction data were processed by multiple auto-processing (including Xia2 and Autoproc) pipelines on ISPYB^60–62^. The high-resolution cutoff was determined by CC ½ above 0.3. The resolution values used in the analysis were the results of the Xia2 processing pipeline. Datasets were truncated using Aimless in CCP4-i suite^63^. Phasing of RECQL5 was done by Phaser^64^, using the D1 and D2 domains of RECQCL5 (PDB id 5LB8) as search models. The search model for phasing the nanobody was chain A of 2X1O. Further refinement was done by COOT and Refmac5 in CCP4-i suite ^65, 66^.

## Data availability

The structures of RECQL5 in complex with engineered nanobodies are deposited in the Protein Data Bank (PDB), with PDB IDs listed as follows.

| Nanobody/Gluebody involved | PDB ID |
| --- | --- |
| G1-001 | 7ZML |
| G2-001 | 7ZMM |
| G3-048 | 7ZMN |
| G3-052 | 7ZMO |
| G3-055 | 7ZMP |
| G2*-006 | 7ZMQ |
| G2*-011 | 7ZMR |
| G4-043 | 7ZMS |
| G5-006 | 7ZMT |
| G5-006 | 7ZMV |

The PDB IDs of structures used in the nanobody interface analysis are listed as follows - 2X1O, 2X1P, 2X1Q, 2X6M, 2X89, 2XT1, 2XV6, 2XXC, 2XXM, 3CFI, 3DWT, 3EAK, 3EBA, 3EZJ, 3G9A, 3K1K, 3K74, 3OGO, 3P0G, 3SN6, 3STB, 3ZKQ, 3ZKS, 4AQ1, 4BEL, 4BFB, 4C57, 4C58, 4C59, 4CDG, 4EIG, 4EIZ, 4EJ1, 4FHB, 4GFT, 4GRW, 4HEM, 4HEP, 4I0C, 4I13, 4I1N, 4IOS, 4JVP, 4KML, 4KRL, 4KRM, 4KRN, 4KRO, 4KRP, 4LDE, 4LDL, 4LDO, 4LGP, 4LGR, 4LGS, 4LHJ, 4LHQ, 4MQS, 4MQT, 4N9O, 4OCL, 4OCM, 4OCN, 4P2C, 4QGY, 4QKX, 4QLR, 4QO1, 4S10, 4S11, 4W6W, 4W6X, 4W6Y, 4WEM, 4WEN, 4WEU, 4WGV, 4WGW, 4X7C, 4X7D, 4X7E, 4X7F, 4XT1, 4Y8D, 4Z9K, 4ZG1, 5BOP, 5BOZ, 5C1M, 5C2U, 5C3L, 5DA0, 5DA4, 5DFZ, 5DXW, 5000, 5E0Q, 5E5M, 5E7B, 5E7F, 5F1K, 5F1O, 5F21, 5F7K, 5F7L, 5F7M, 5F7N, 5F7W, 5F7Y, 5F8Q, 5F8R, 5F93, 5F97, 5F9A, 5F9D, 5FWO, 5G5R, 5G5X, 5GXB, 5H8D, 5H8O, 5HDO, 5HGG, 5HVF, 5HVG, 5HVH, 5IMK, 5IML, 5IMM, 5IMO, 5IP4, 5IVN, 5IVO, 5JA8, 5JA9, 5JDS, 5JQH, 5LHN, 5LHP, 5LHQ, 5LHR, 5LMJ, 5LMW, 5LZ0, 5M13, 5M14, 5M15, 5M2M, 5M2W, 5M30, 5M7Q, 5M94, 5M95, 5MJE, 5MP2, 5MWN, 5MY6, 5MZV, 5NBD, 5NBL, 5NBM, 5NLU, 5NLW, 5NM0, 5NML, 5NQW, 5O02, 5O03, 5O04, 5O05, 5O0W, 5O2U, 5O8F, 5OCL, 5OJM, 5OMM, 5OMN, 5OVW, 5TD8, 5TJW, 5TOJ, 5TOK, 5TP3, 5U64, 5U65, 5UK4, 5UKB, 5USF, 5VAK, 5VAN, 5VL2, 5VLV, 5VM0, 5VM4, 5VM6, 5VNV, 5VNW, 5WB1, 5WB2, 5Y7Z, 5Y80, 6B20, 6B73, 6C5W, 6C9W, 6DBA, 6DBD, 6DBE, 6DBF, 6DBG, 6EHG, 6EQI, 6EY0, 6EY6, 6EZW, 6F0D, 6FE4, 6FUZ, 6FV0, 6GZP, 6H02, 6H15, 6H16, 6H1F, 6H6Y, 6H6Z, 6H70, 6H71, 6H72, 6H7J, 6H7L, 6H7M, 6H7N, 6H7O, 6I6J, 6IBL, 6MXT, 6QTL, 6R7T, 6O3C, 6OS0, 6OS1, 6OS2, 6OYH, 6OYZ, 6OZ6, 6Q6Z, 6QD6, 6QGW, 6QGX, 6QGY, 6QPG, 6QUZ, 6QV0, 6QV1, 6QV2, 6QX4, 6RNK, 6RTW, 6RTY, 6RU3, 6RU5, 6RUL, 6RUM, 6RUV, 6RVC, 6S0Y, 6SGE, 6SSI, 6SSP, 6TEJ, 6TYL, 6SWR, 6U12, 6U14, 6U50, 6U51, 6U52, 6U53, 6U54, 6U55, 6VI4, 6WAQ, 6WAR, 4DK3, 4DK6, 4DKA, 4KDT, 4KSD, 4PIR, 5J1S, 5J1T, 5M2I, 5M2J, 6GJQ, 6GJS, 6GJU, 6GK4, 6GKD, 6GWN, 6GWP, 6GWQ, 6HD8, 6HD9, 6HDA, 6HDB, 6HDC, 6HEQ, 6HER, 6HHD, 6HHU, 6HJX, 6HJY, 6I2G, 6I8G, 6I8H, 6IBB, 6ITP, 6ITQ, 6JB2, 6JB5, 6JB8, 6JB9, 6DO1, 6DYX, 6F2G, 6F2W, 6FPV, 6GCI, 6GS1, 6GS4, 6GS7, 6N4Y, 6N50, 6NFJ.

## Acknowledgements

We acknowledge the support and use of resources of Instruct-ERIC (PID6873), part of the European Strategy Forum on Research Infrastructure (ESFRI), and the Research Foundation – Flanders (FWO) for their support to the Nanobody discovery. Next Generation Chemistry at the Rosalind Franklin Institute is supported by the EPSRC (EP/V011359/1). We acknowledge the efforts of Katleen Willibal for the technical assistance during the discovery of wild-type nanobodies we used in this study. The crystallographic screen was supported by the XChem facility at Diamond Light Source (proposal ID LB26998). We acknowledge the support from all the staff of Diamond Light Source. The SGC was a registered charity (number 1097737) that received funds from AbbVie, Bayer Pharma AG, Boehringer Ingelheim, Canada Foundation for Innovation, Eshelman Institute for Innovation, Genome Canada, Innovative Medicines Initiative (EU/EFPIA) [ULTRA-DD grant no. 115766], Janssen, Merck KGaA Darmstadt Germany, MSD, Novartis Pharma AG, Ontario Ministry of Economic Development and Innovation, Pfizer, São Paulo Research Foundation-FAPESP, Takeda, and Wellcome [106169/ZZ14/Z]. M.Y. was on the China Scholarship Council – Nuffield Department of Medicine (CSC-NDM) award at the University of Oxford. G.Y. was a Clarendon Scholar at the University of Oxford. G.Y. and R.J.C.G. were supported by the Calleva Research Centre for Evolution and Human Sciences at Magdalen College, Oxford.

## Author contributions

Conceptualization: M.Y., F.V.D.

Methodology: M.Y., M.M. M.F., E.M., N.D., H.A.

Investigation: M.Y., M.M. M.F., G.Y., H.L., D.M.

Visualization: M.Y. and J.A.N.

Funding acquisition: F.V.D., O.G., K.D., R.J.C.G. and B.G.D.

Project administration: F.V.D.

Supervision: F.V.D., O.G. and B.G.D.

Writing – original draft: M.Y.

Writing – review & editing: M.Y., M.M, J.A.N., M.F., E.M., N.D.W., L.K., A.T., G.A.B., G.Y., H.L., V.L.R., D.M., B.G.D., R.J.C.G., K.D., O.G. and F.V.D.

## Extended Data figures

**Extended Data Figure 1.**
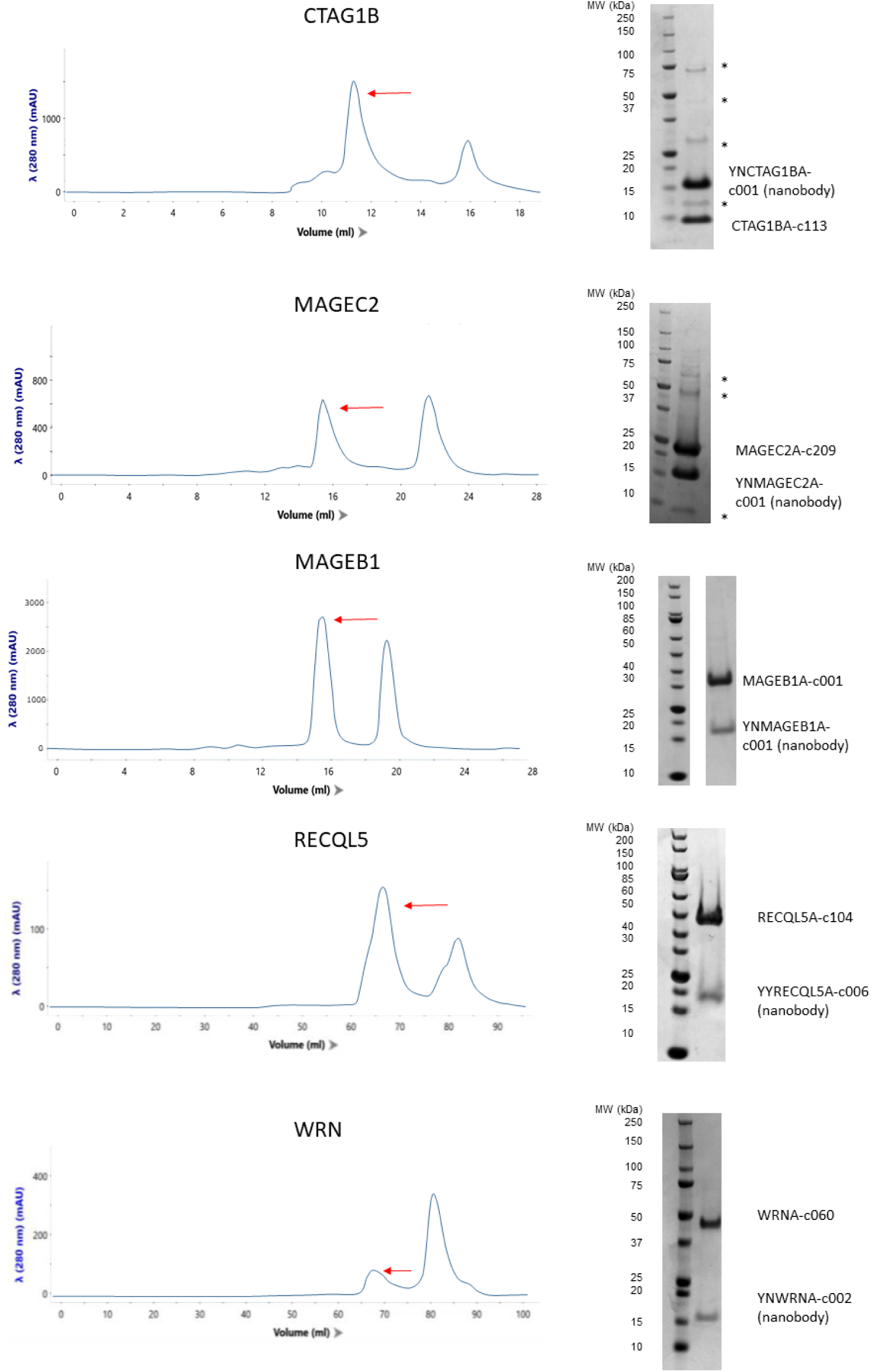
Size exclusion chromatography profiles (left) and SDS-PAGE images (right) for each target protein. The red arrows show the protein peak fractions collected for crystallization trials. ‘*’ indicates contaminants in the protein samples.

**Extended Data Figure 2.**
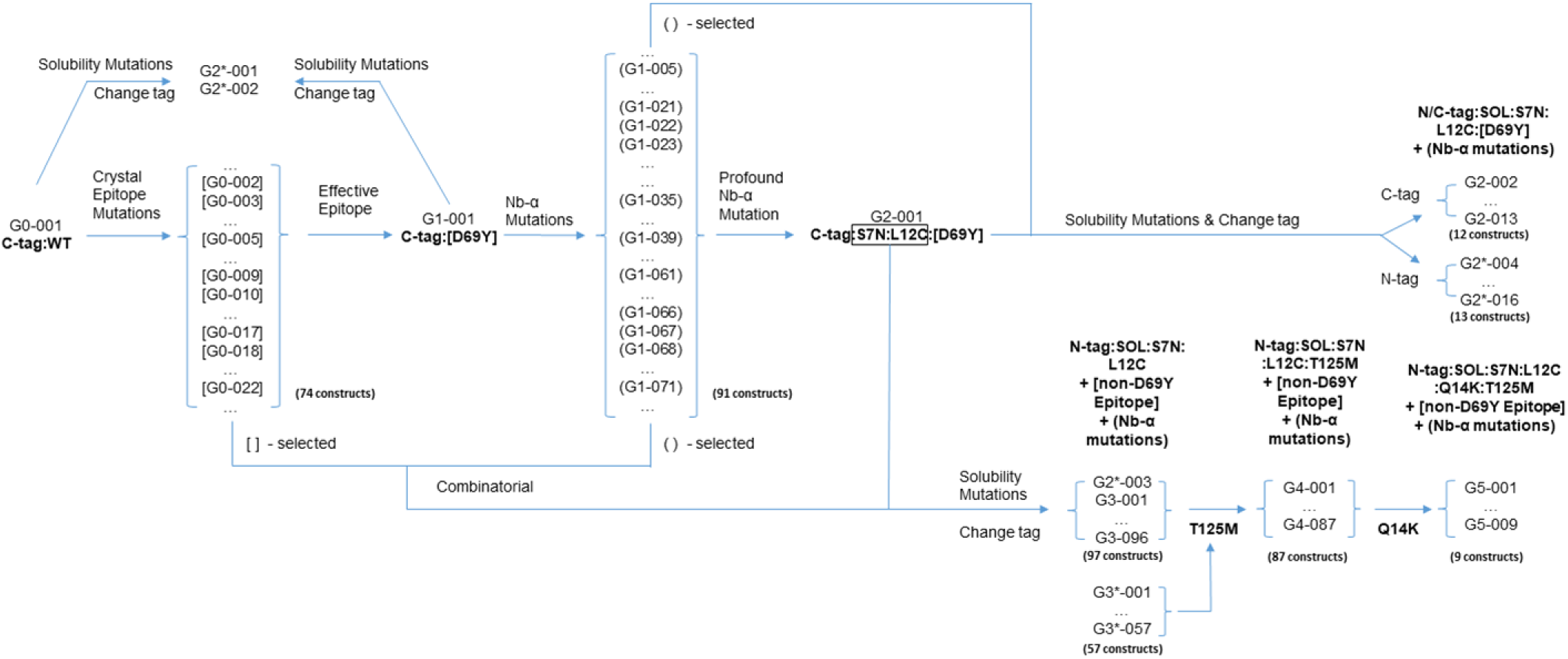
The road map of the several iterations of mutagenesis on RECQL5 nanobody scaffold. Variants in ‘[]’ are selected crystal epitope mutants and in ‘()’ are selected Nb-α mutants for subsequent iterations of mutagenesis.

**Extended Data Figure 3.**
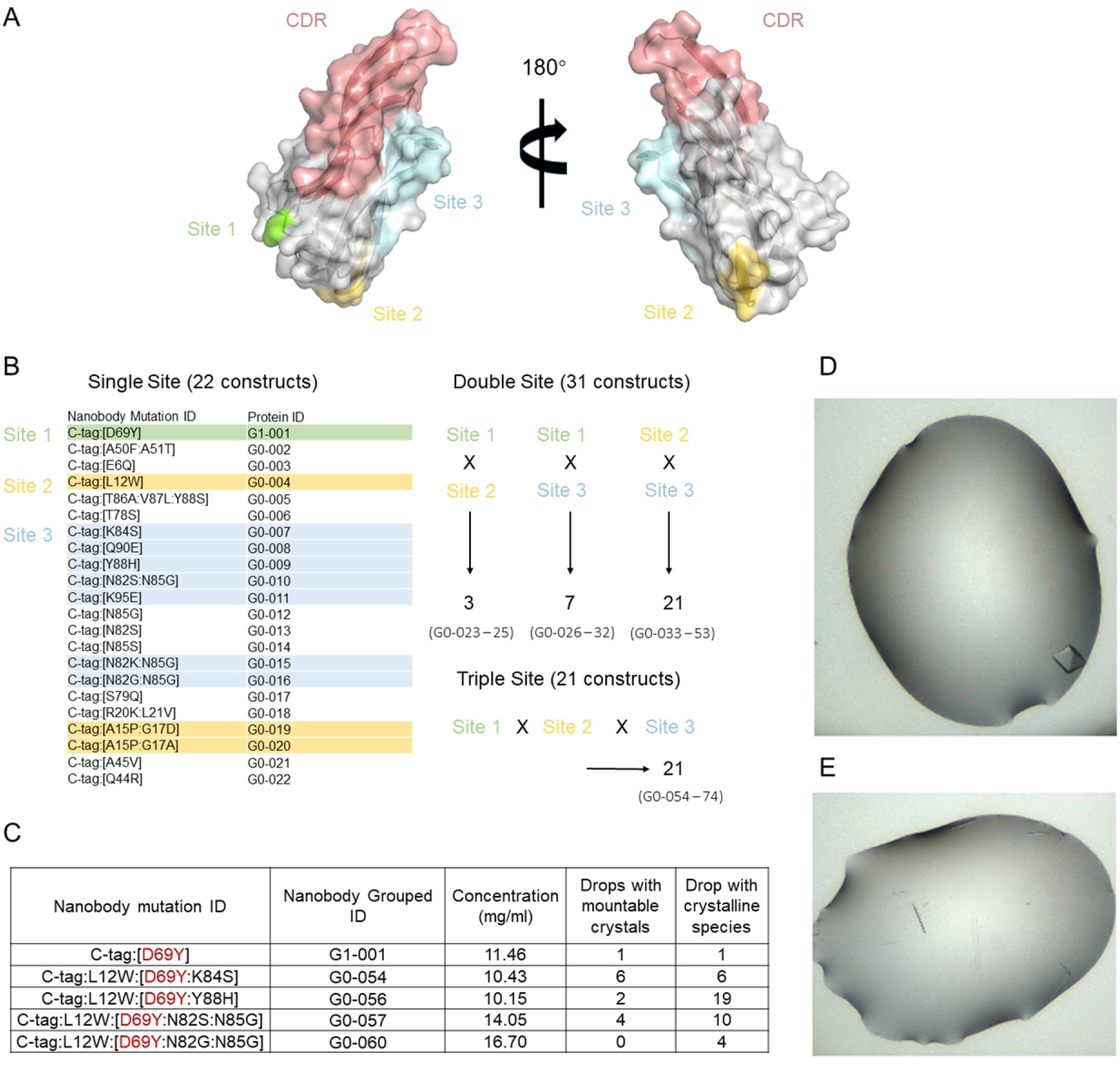
The crystal epitope mutation D69Y on the scaffold rescues the RECQL5 nanobody to be effective again as a crystal chaperone. A) The crystal epitope sites on the surface of a RECQL5 nanobody (C-tag:WT) are coloured in green, yellow and cyan. CDR regions are coloured in light brown. B) The combinatorial mutation designs. 22 single site mutations were chosen from the list of mutations provided by the crystal epitope server. 11 of the single site mutations were then selected to generate double-site and triple-site mutations thereafter, which resulted in 74 nanobody variants for crystallization trials. Colours are consistent with the sub-figure A. C) The crystal epitope variants that yielded crystals in the Hampton Index 3 Screen. All effective nanobody variants shared the D69Y mutation, highlighted in red. D) G1-001 crystal. E) G0-057 crystals.

**Extended Data Figure 4.**
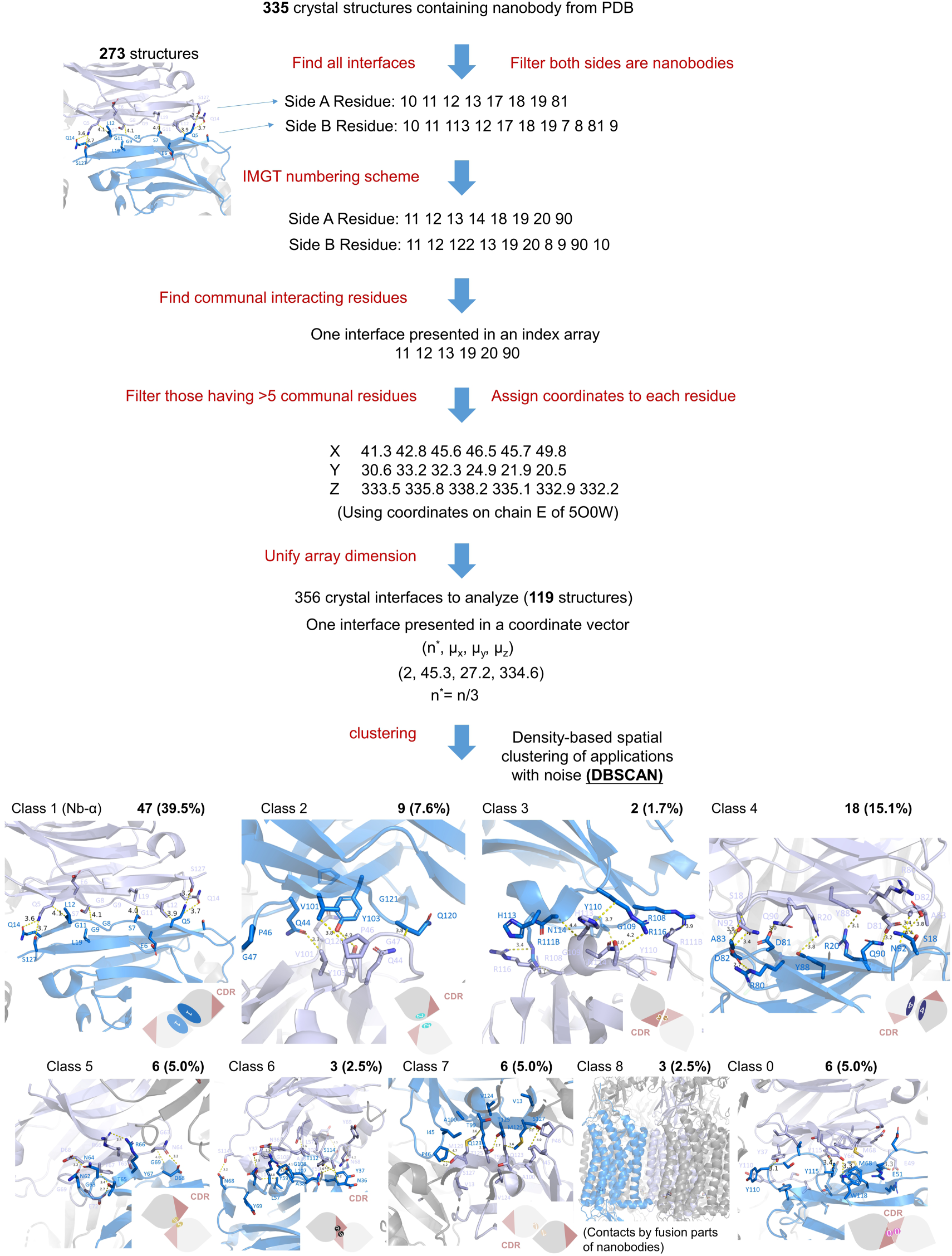
The computational workflow of nanobody-nanobody crystal contact analysis and classification results. The texts in red represent actions taken in each step. Structure cartoons in marine and light blue represent two nanobody molecules participating in the interface. Residues represented as sticks are interacting residues of the current nanobody molecules. Simplified nanobody sketches are shown at the bottom corners, and are consistent with Figure 2. Class 8 interfaces are between fusion parts of the nanobody and therefore are not counted as nanobody-nanobody interfaces.

**Extended Data Figure 5.**
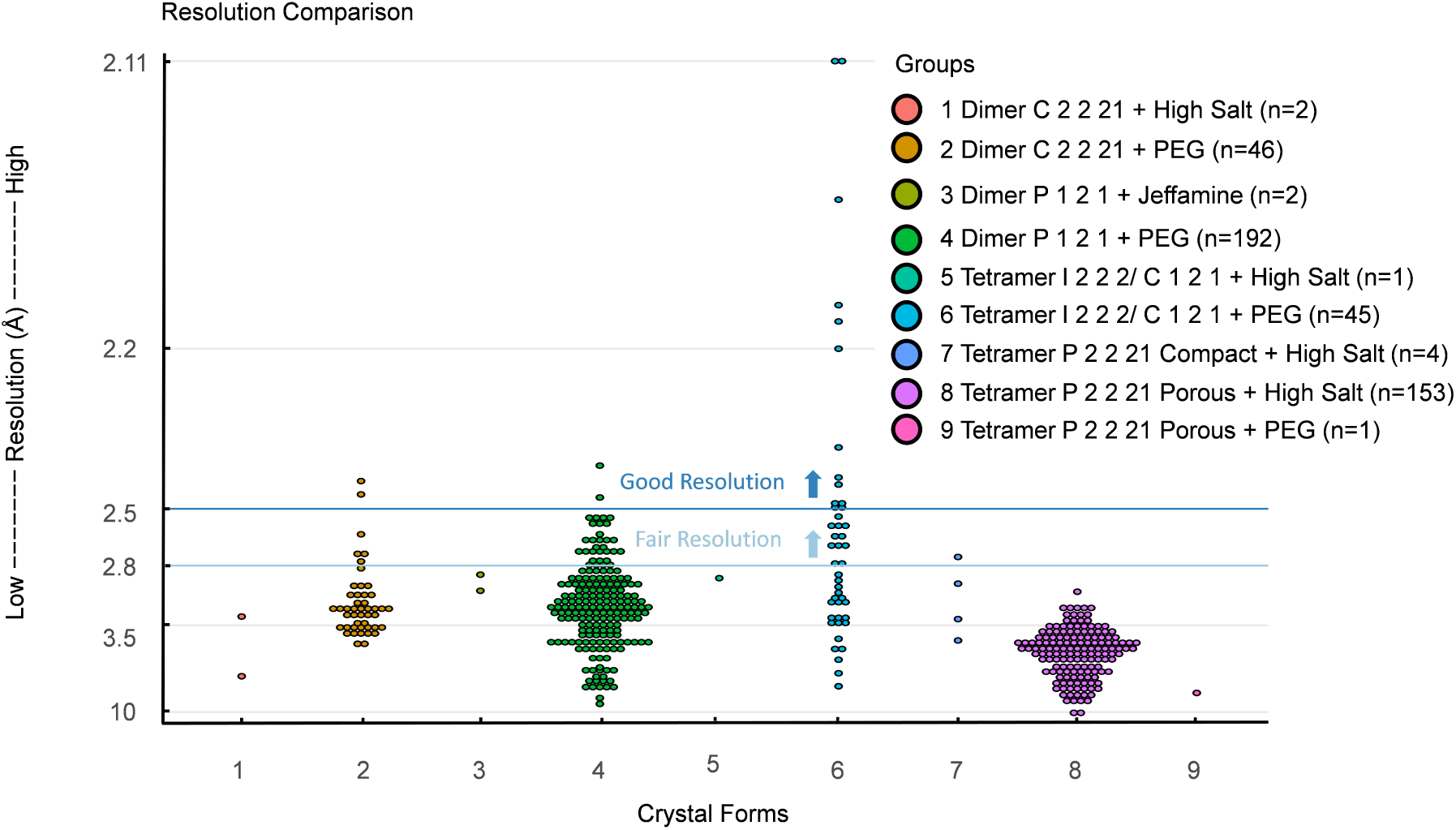
Resolution comparison of different crystal forms from RECQL5:nanobody complexes. Each combination of crystal forms and crystallization condition categories are grouped and indicated in different colors. The crystals are not in fragment-soaking conditions. Resolution indicated here uses the criteria CC ½ > 0.3 from the ISPYB auto-processing pipeline without any further data truncation.

**Extended Data Figure 6.**
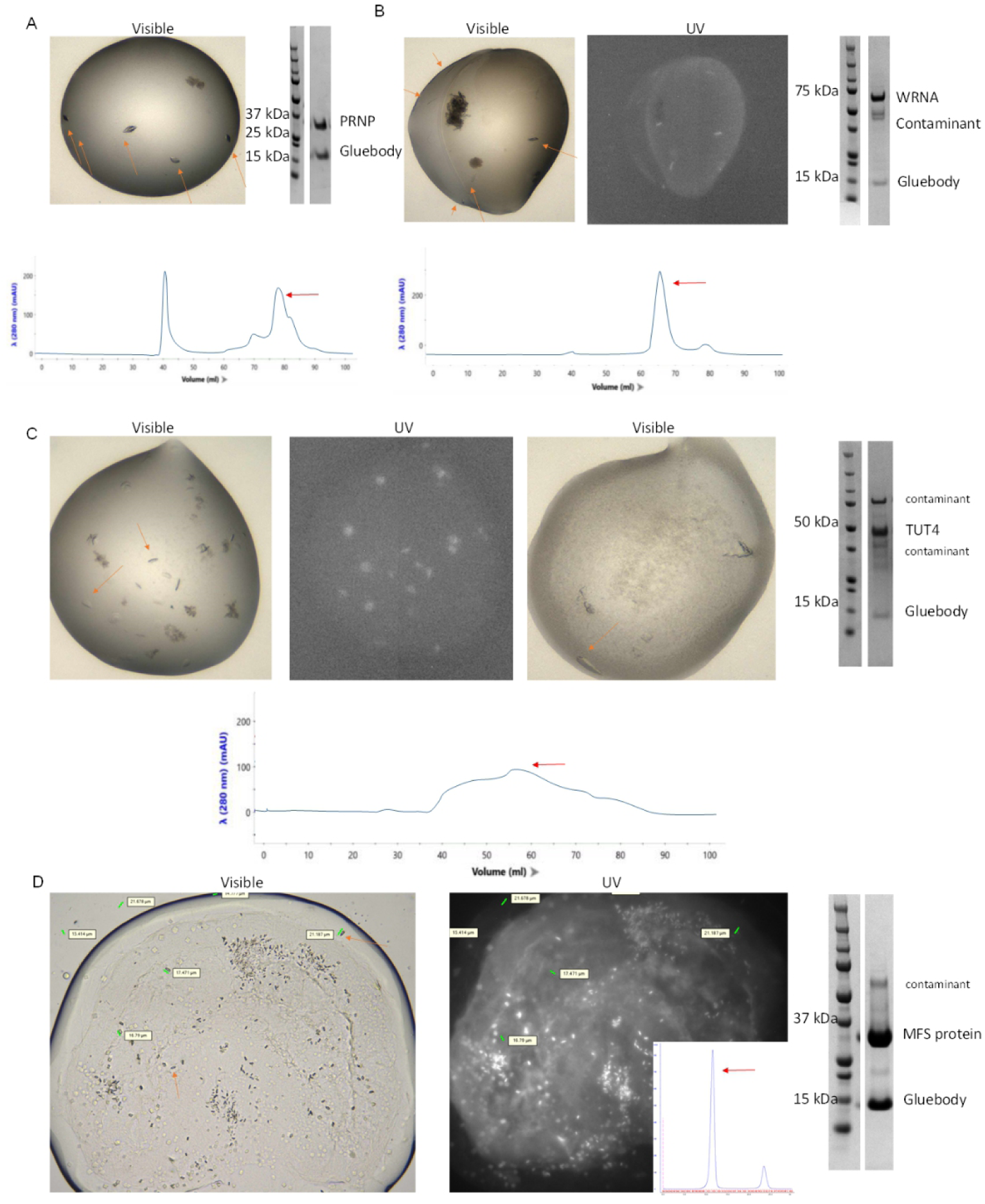
Crystals from PRNP, WRN, TUT4 and MFS protein in complex with respective Tetra-Gluebodies. There are visible and UV images for crystals and SDS-PAGE, SEC profiles showing the quality of the protein. The protein peaks in SEC collected for crystallization trials are indicated by the red arrows. Crystals are indicated by orange arrows. A) PRNP:Gluebody complex crystals. Crystallization happened at 14 mg/ml, 20 °C in the condition of 0.2M Magnesium chloride hexahydrate, 0.1M tris pH 8.5 25% w/v PEG3350, with a protein:precipitant ratio of 1:2. B) WRN:Gluebody complex crystals. Crystallization happened at 27.3 mg/ml, 20 °C in the condition of 0.15 M potassium bromide, 30% w/v polyethylene glycol monomethyl ether 2,000, with a protein:precipitant ratio of 1:2. C) TUT4:Gluebody complex crystals. There are several conditions yielding crystals. In the visible image on the left, the condition is 0.2 M magnesium chloride hexahydrate, 0.1 M BIS-TRIS pH 6.5, 25% w/v polyethylene glycol 3,350, with a protein:precipitant ratio of 1:2. In the visible image on the right, the condition is 0.1 M HEPES pH 7.5, 2.0 M ammonium sulfate, with a protein:precipitant ratio of 1:2. All crystallization happened at 14 mg/ml, 20 °C. D) The LCP crystals formed in the condition of 0.1M NaCl 0.1M Li_2_SO_4_ 40%v/v PEG200 0.1M MES pH 6 at 40 mg/ml, 20 °C.

**Extended Data Figure 7.**
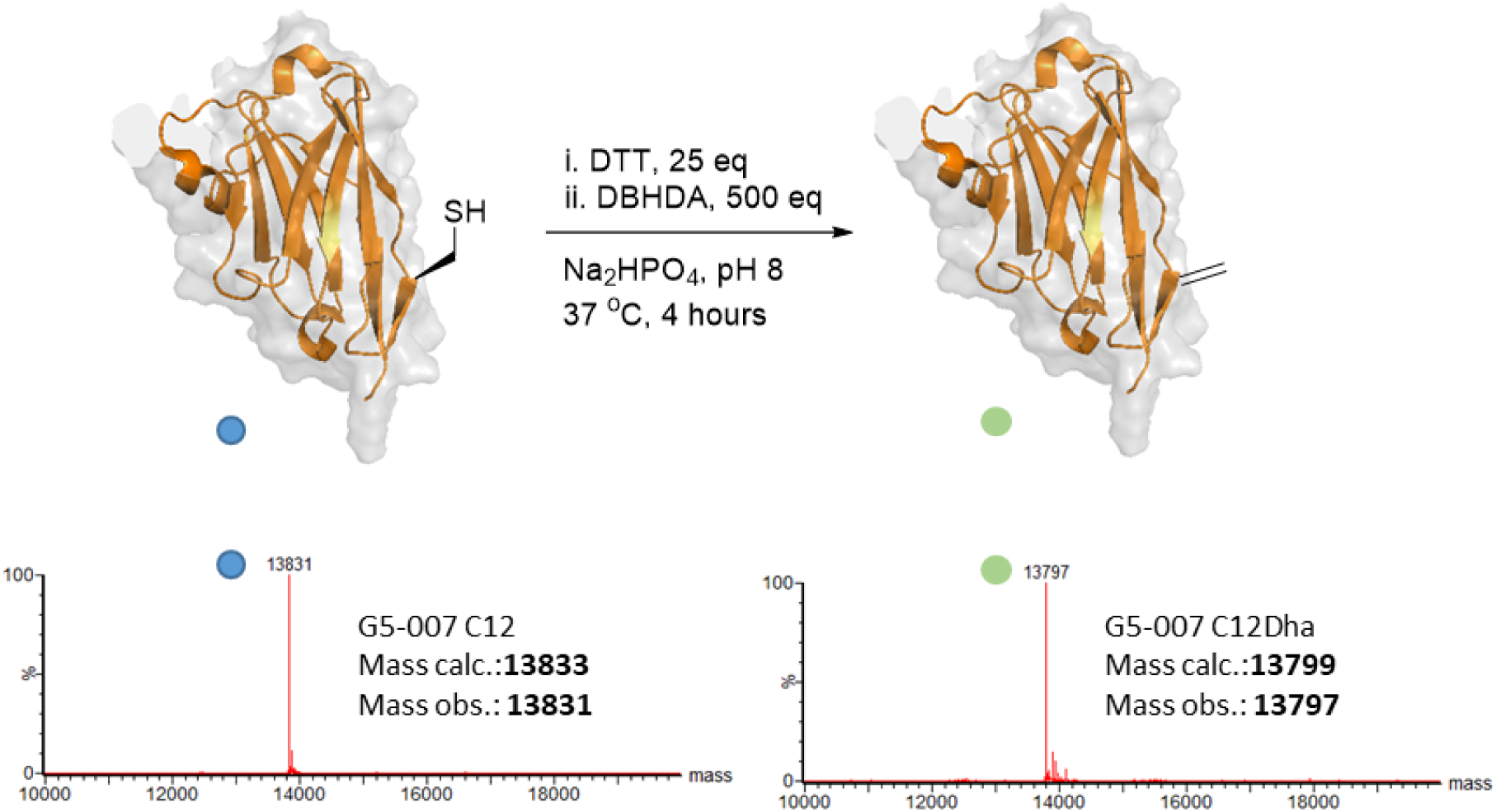
Chemical conversion of the Cys12 on G5-007 nanobody to dehydroalanine (Dha). To a solution of G5-007 nanobody (1 mL, 1 mg/mL, 72 μM) in Na_2_HPO_4_ buffer (50 mM, pH 8.0), 0.28 mg (25 eq.) of DTT were added. The solution was shaken at room temperature for 20 min and then the protein was treated with 22 μL of 0.5M DBHDA (2,5-dibromohexanediamide) in DMSO (500 eq.) and heated to 37 °C for 4 h, at which point analysis by mass spectrometry showed reaction completion. The protein was purified using a PD Miditrap™ G-25 column (Cytiva #28918008) pre-equilibrated with Na_2_HPO_4_ buffer (50 mM, pH 8.0).

**Extended Data Figure 8.**
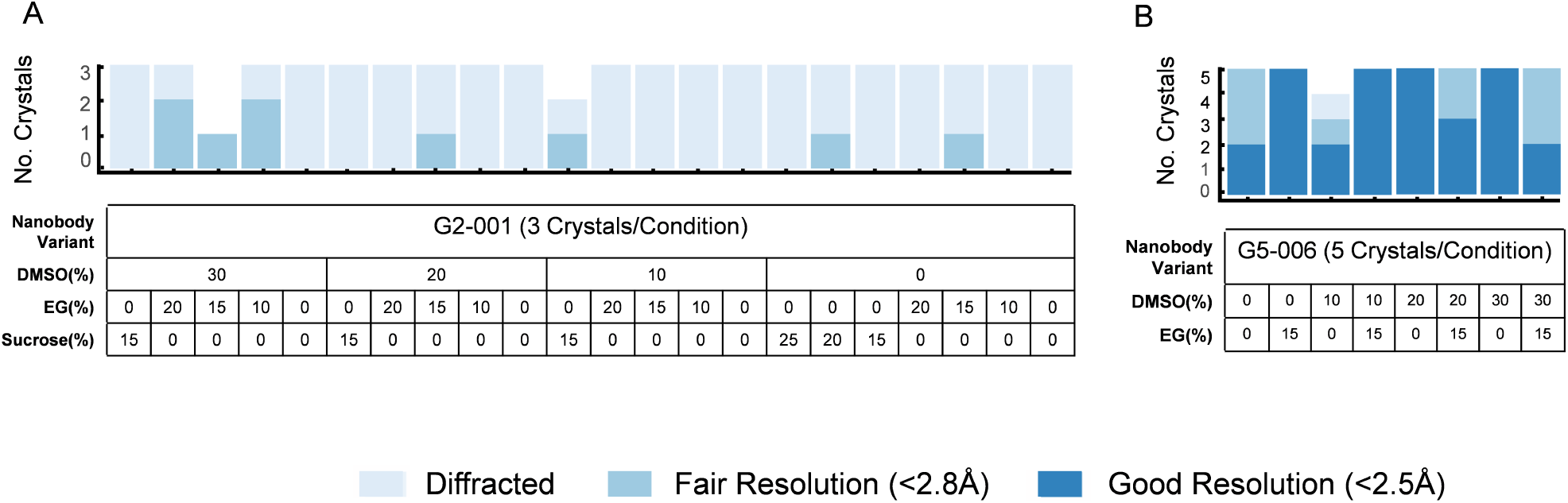
Diffraction quality of G5-006 (A) and G5-009 (B) soaked in different cryo-protectants, shaded according to diffraction resolution. In the bar plot, deep blue fractions represent good resolution (<2.5 Å) crystal percentage, blue bar heights represent fair resolution (<2.8 Å) crystal percentage and the total heights of the bars represent percentage of crystals that diffracted. The numbers of crystals, nanobody variants and soaking conditions in each group are indicated in the table below each bar.

**Extended Data Figure 9.**
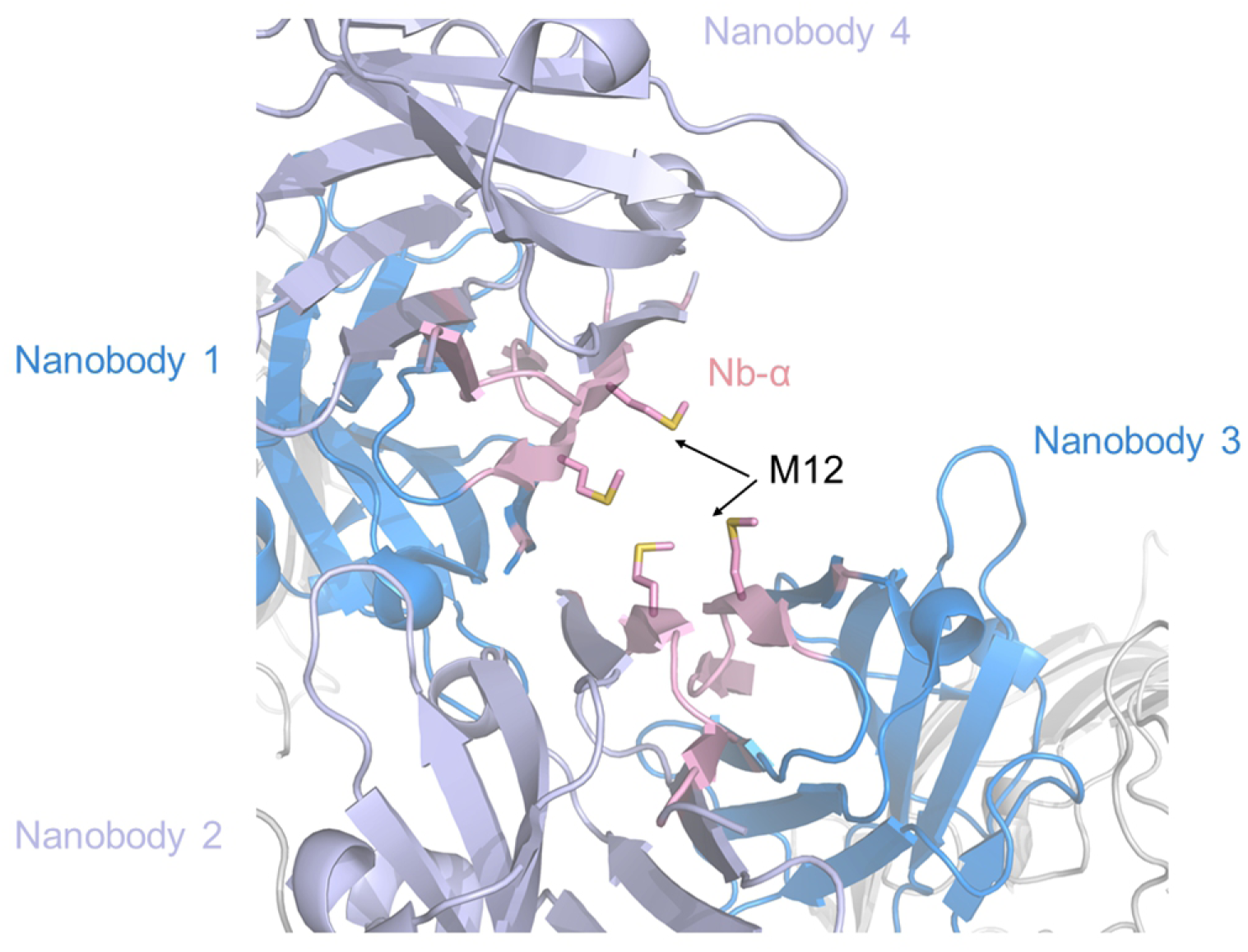
Tilted back-to-back packing of nanobody dimers from PDB id 6QGY. The C12 is replaced by M12, resulting in the tilted packing. Nb-α is also present in this crystal lattice. Molecules colored in marine and light blue are nanobody molecules. Nb-α is colored pink. The target protein molecules of the nanobodies are colored grey.

**Extended Data Figure 10.**
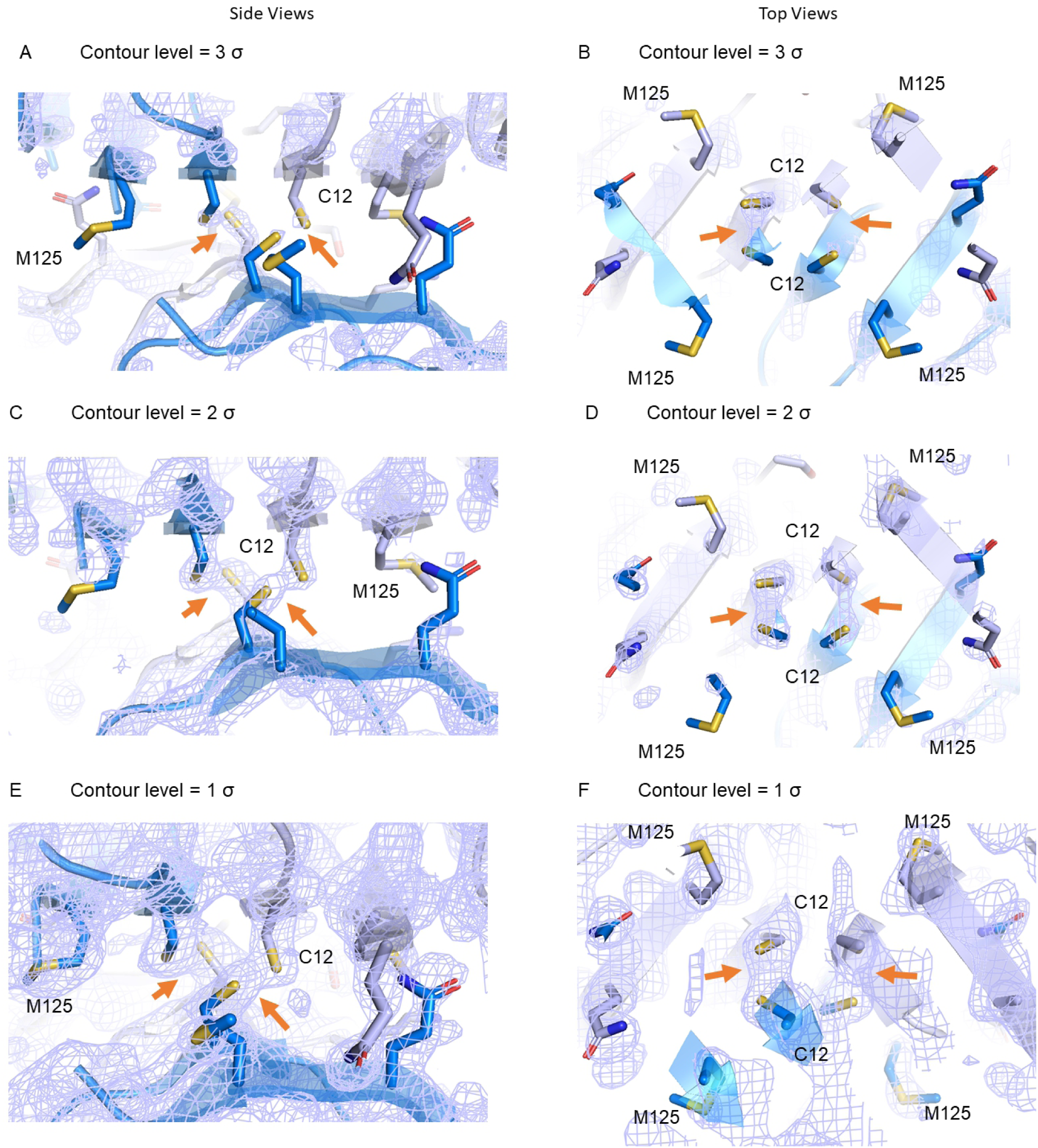
Electron density (pointed to by yellow arrows) implies possible bonds between the sulfur atoms in the C-C interface core. C12 and M125 are presented in stick and the sulfur atoms are colored yellow. The electron density around C12 is indicated in light blue mesh with the contour levels at 1σ, 2σ and 3σ, viewed from the top and the side.

## Extended Data tables

**<u>Extended Data Table 1</u>** The crystallization experiment on a selection of target:nanobody pairs with naïve nanobody scaffolds.

**<u>Extended Data Table 2</u>** Detailed DBSCAN result with full list of PDB codes of each nanobody-nanobody interface category. In the ’Pattern name’ column, first four letters represent PDB and the digit after ’_’ represents the No. of the pattern in the structure. In the ‘Pattern’ column, each number represents the IMGT normalized index of a residue of the nanobody scaffold involved in the pattern.

**<u>Extended Data Table 3</u>** Full list and the crystallization result of nanobody variants tested in the RECQL5:nanobody system

**<u>Extended Data Table 4</u>** Full list of diffraction data collected from crystals of all nanobody variants tested in the RECQL5:nanobody system. Resolution indicated here uses the criteria CC ½ > 0.3 from the ISPYB auto-processing pipeline without any further data truncation.

**<u>Extended Data Table 5</u>** Detailed information with numeric values of each cell in Figure 5A

